# Pathogen Evolution when Transmission and Virulence are Stochastic

**DOI:** 10.1101/584425

**Authors:** Pooya Aavani, Sean H. Rice

## Abstract

Evolutionary processes are inherently stochastic, since we can never know with certainty exactly how many descendants an individual will leave, or what the phenotypes of those descendants will be. Despite this, models of pathogen evolution have nearly all been deterministic, treating values such as transmission and virulence as parameters that can be known ahead of time. We present a broadly applicable analytic approach for modeling pathogen evolution in which vital parameters such as transmission and virulence are treated as random variables, rather than as fixed values. Starting from a general stochastic model of evolution, we derive specific equations for the evolution of transmission and virulence, and then apply these to a particular special case; the SIR model of pathogen dynamics. We show that adding stochasticity introduces new directional components to pathogen evolution. In particular, two kinds of covariation between traits emerge as important: covariance across the population (what is usually measured), and covariance between random variables within an individual. We show that these different kinds of trait covariation can be of opposite sign and contribute to evolution in very different ways. In particular, probability covariation between random variables within an individual is sometimes a better way to capture evolutionarily important tradeoffs than is covariation across a population. We further show that stochasticity can influence pathogen evolution through directional stochastic effects, which results from the inevitable covariance between individual fitness and mean population fitness.

## Introduction

All organisms live in environments that are, to some degree, unpredictable, and this unpredictability influences both an individual’s reproductive success and the phenotype of its offspring (through environmental effects on development). This uncertainty is likely to be pronounced for pathogens, which could be subject to both stochastic uncertainty within a host (such as variation in the host’s immune response), and in the host’s environment. Stochastic fitness has been shown to impact evolutionary dynamics in ways that are not obvious from deterministic models [1, 2, 3].

One well studied process that will create stochasticity in transmission rate is variation in interactions between susceptible and infected hosts. In many infectious disease models, the population is assumed to be large and evenly mixed, meaning that every individual contacts others at a fixed rate determined by their frequency [4]. In reality, however, infected individuals often have limited and variable numbers of contacts with susceptible individuals [5]. For example, some epidemiological studies have shown that during disease spreading, the population is heterogenous, with a few particularly individuals responsible for the majority of transmission events [6, 7, 8, 9, 10]. In particular Woolhouse et. al identified an empirical relationship that in many disease systems, the most infectious 20% of individuals are responsible for 80% of the total infections [11]. Stochastic variation in transmission thus appears to be a common feature of pathogen systems.

Another important component of pathogen fitness, virulence, is also likely to be stochastic, in part because of variation in the internal environment of the host – particularly the hosts immune system. For example, Brodin and Davis document striking variation of individual immune response in humans [12]. This translates into variation in virulence rates for a particular pathogen, with virulence being relatively low in some hosts and high in others. Even within a host, variation in virulence could be a result of uncertainty in the ability of pathogens to evade the immune system and induce pathogenic effects.

Variation in transmission and virulence has also been seen in other studies, with the further observation that the pattern of variation may itself be influenced by external environmental variables that the hosts experience. For example, in an experimental study of *Daphnia magna* infected by the bacterial parasite *Pasturia ramosa*, it was shown that changing environmental factors, such as nutrient availability or temperature, can alter the variance in both virulence and transmission, as well as the covariance between them [13, 14, 15].

Figure (1) shows the distribution of mortality rates as a function of time since infection (an estimator of 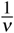) for human immunodeficiency virus 1 (HIV1) [data from [16]]. This data suggests both a high variance in virulence, and that the distribution of virulence values is asymmetrical.

**Figure 1:**
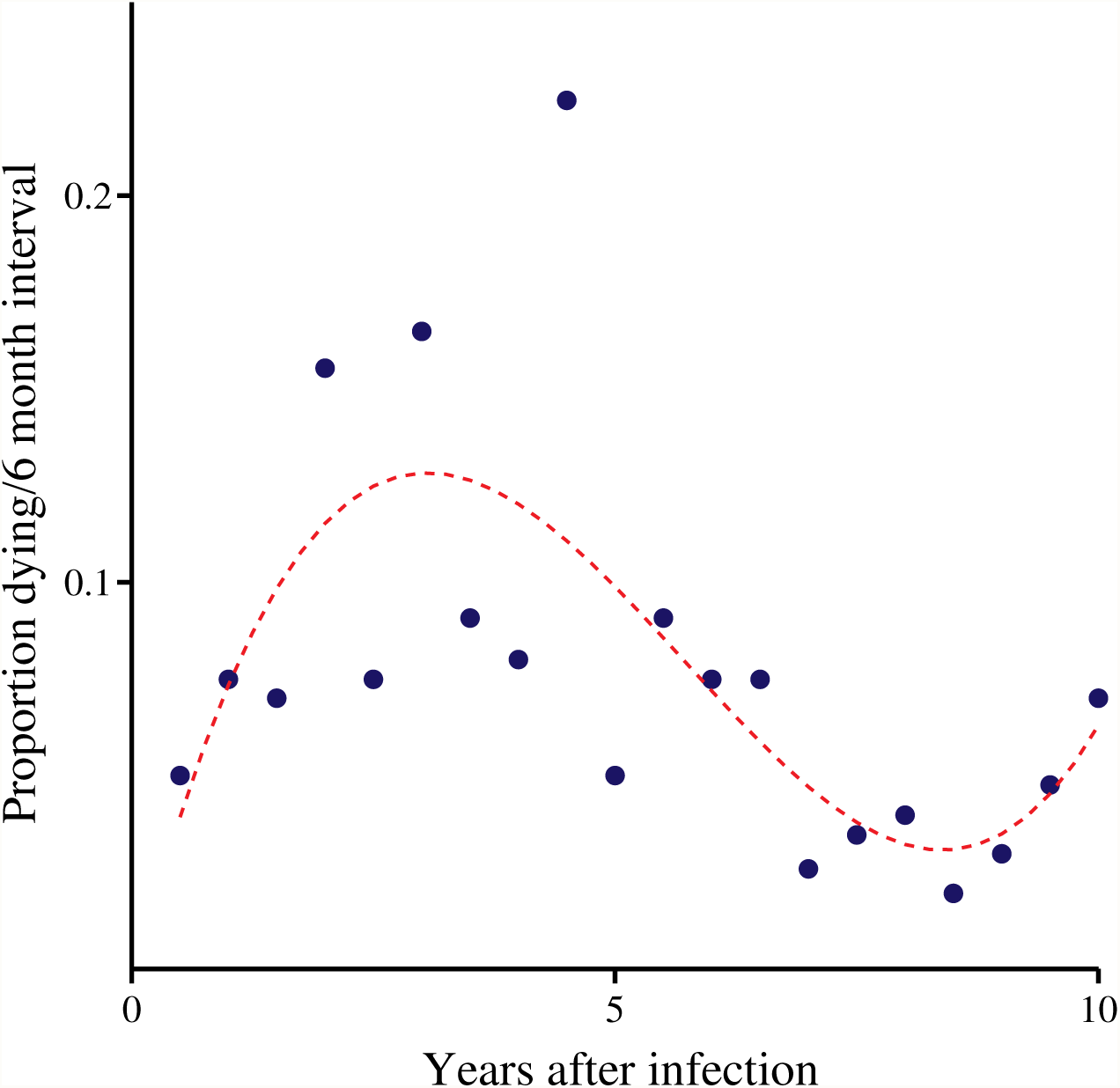
Distribution of mortality/year (virulence) during 10 consecutive years after infection with HIV1. The red fitted third-degree polynomial shows the positive skewness of data and so the high variance in virulence. The data are taken from [16], that conducted the cohort study of patients after infection with HIV1 and free of clinical acquired immunodeficiency syndrome(AIDS).

In a stochastic evolutionary model, we treat both the number of offspring that each individual will produce and the phenotype of those offspring, as random variables – having distributions of possible values. Saying that something is a “random variable” does not mean that it is completely random in the sense that we can make no predictions about it – only that there is some uncertainty about the exact value that it will ultimately take. Therefore, when we treat transmission as a random variable, it means only that when a pathogen infects a host, we can not predict with certainty the exact number of subsequent hosts that will be infected through transmission from the current host. Instead, we work with the probability distribution of possible transmission events. Similarly, saying that virulence is stochastic just means that we can not predict exactly how much harm a newly acquired pathogen will ultimately do to its host.

A random variable has a distribution, and we can, in principle, predict the mean, variance and other moments of this distribution. For evolution with stochastic selection, we therefore calculate the expected (probability mean) change in the (frequency) mean phenotype. In the absence of migration, this is given by the stochastic Price equation [2]:

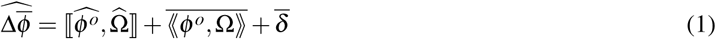

In Equation (1), *ϕ*^*o*^ denotes the mean phenotype of an individual’s offspring (or of that individual in the future), and Ω denotes relative fitness – individual fitness divided by mean population fitness (see Table 1). 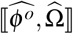 is the covariance, across the population, between expected offspring phenotype and expected relative fitness. We refer to this as the frequency covariance, since it involves the frequencies of individuals in the population. Because we are now treating *ϕ*^*o*^ and Ω as random variables, however, each individual has a probability distribution of possible values of offspring phenotype and relative fitness. These values can thus covary within an individual, producing a probability covariance that we denote ⟪*ϕ*^*o*^, Ω ⟫. Figure (2) shows schematic representation of frequency and probability covariances.

**Figure 2:**
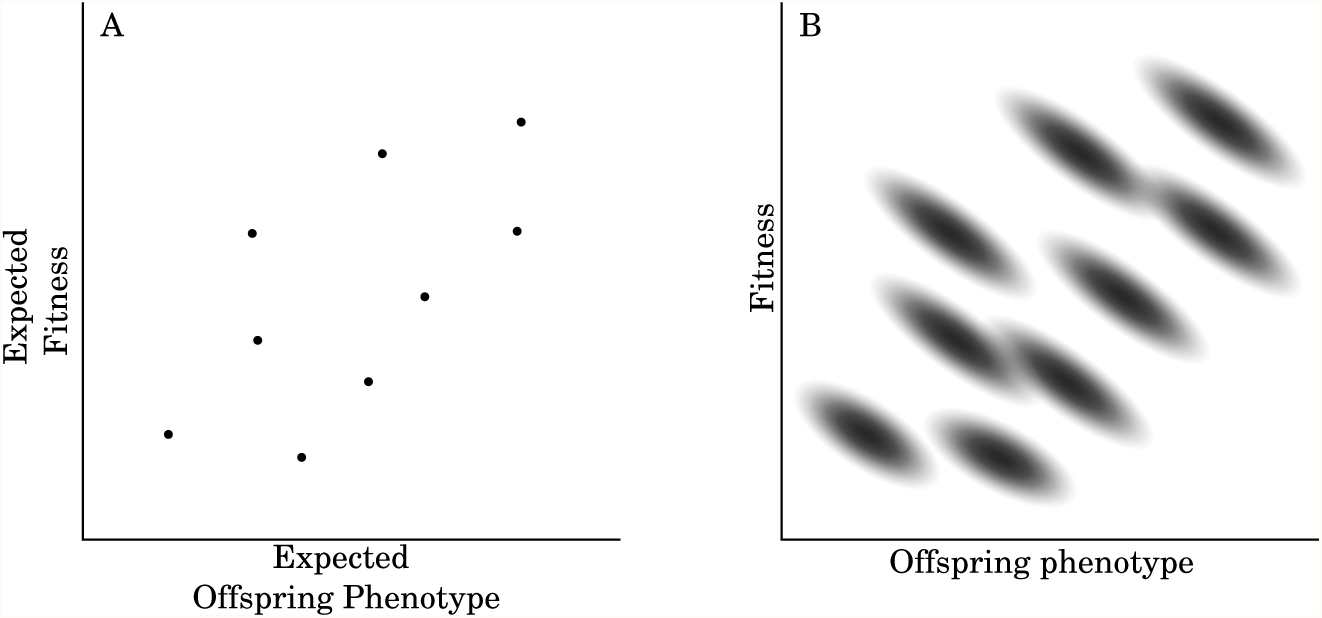
Frequency covariance versus Probability covariance. Figure (A) shows the relation between expected offspring phenotype and expected fitness, 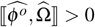. Figure (B) shows that each individual has a probability distribution of possible values of offspring phenotype and relative fitness, 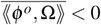.

**Table 1:** Symbols and Notation

| Symbol | Meaning |
| --- | --- |
| $\bar{X}$ | Average value of X across its frequency distribution |
| $\hat{A}$ or $E(A)$ | Expected value of random variable A |
| $A^*$ | Difference between random variable A and its expectation $\hat{A}$ (i.e., $A - \hat{A}$) |
| $\llbracket^2 X\rrbracket$ | Variance in the value of X, across its frequency distribution |
| $\llbracket^j X\rrbracket$ | $j^{\text{th}}$ central moment of the frequency distribution of X |
| $\langle\langle^j A\rangle\rangle$ | $j^{\text{th}}$ central moment of the probability distribution of random variable A |
| $\llbracket X, Y\rrbracket$ | Frequency covariance of X and Y |
| $\langle\langle A, B\rangle\rangle$ | Probability covariance of random variables A and B |
| $w$ | Absolute fitness (number of descendants after one generation) |
| $\Omega$ | Relative fitness ($\Omega = \frac{w}{\bar{w}}$), conditional on $\bar{w} \neq 0$ |
| $\phi$ | Phenotype of an individual |
| $\phi^o$ | Average phenotype of an individual's offspring |
| $\delta$ | $\phi^o - \phi$ |
| $\theta$ | Birth rate of susceptible hosts |
| $d$ | Death rate of the host in the absence of infection (Background Mortality) |
| $c$ | The rate that infected groups become immune to further infections (Recovery Rate) |
| $\beta_i$ | The rate at which susceptible hosts become infected by individuals in $i^{\text{th}}$ infected group (transmission rate) |
| $v_i$ | The induced death rate of the individuals in $i^{\text{th}}$ infected group by the pathogens (virulence rate) |

To illustrate the difference between these two kinds of covariance: Saying that 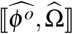 is positive for a population means that individuals within the population that have large values of 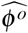 also tend to have large values of 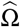. By contrast, saying that ⟪*ϕ*, Ω ⟫ is positive for an individual means that if an individual’s value of *ϕ*^*0*^ ends up being larger than its expected value 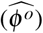, then its value of Ω is also likely to be larger than its expected value 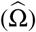. The distinction between probability and frequency operations is critical in modeling stochastic processes in populations. The term *δ* captures processes, such as mutation, that change phenotype during the process of reproduction.

Equation (1) applies to any phenotype. It assumes only that the population is closed, meaning that there is no immigration or emigration. For the case of pathogen evolution, we will consider two phenotypic traits, transmission (*β*) and virulence (*ν*).

In the following section we start by deriving a general stochastic model of evolution and then derive specific equations for evolution of transmission and virulence. We will show that adding stochasticity does not just add noise to our results, it introduces new directional components to pathogen evolution.

### The General Case

Fitness enters into Equation (1) in the form of 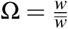, or relative fitness. In order to study selection, we need to write our equations in terms of individual fitness, *w*. This is easy in deterministic models – since 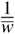 is a constant that we can factor out – but is challenging in stochastic models, in which both *w* and 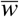 are random variables that are correlated with one another. The solution is to approximate 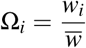 using a Taylor expansion [2, 17] (see Methods and Appendix). Using this approach, we can rewrite the stochastic Price Equation (1) as:

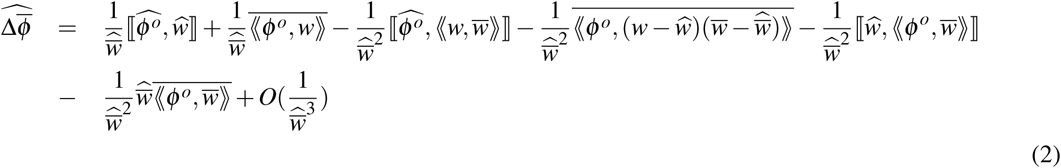

Equation (2) is the general equation of change in the expected mean phenotype when descendant phenotype, and parent fitness are random variables. Often, the “descendants” are assumed to be offspring, but *ϕ*^*o*^ can also measure an individual’s phenotype at a later time, or the phenotype of descendants after many generations. To apply this equation to transmission and virulence of pathogens, we consider a pathogen that has just infected a host, and interpret *ϕ*^*o*^ as the phenotype after that pathogen strain has grown within the host to the point at which it can be transmitted or kill the host.

### Evolution of Transmission and Virulence

We assume that there are *n* pathogen strains, and define *β*_*i*_ as the number of other hosts that will be directly infected by the descendants of strain *i* (*i* = 1, …, *n*), and *ν*_*i*_ as the probability that strain *i* kills its host before transmission. Both *β* and *ν* are random variables, since the exact transmission rate or time until death of the host can not be specified exactly beforehand.

Substituting *β* and *ν* for *ϕ*^*o*^ in Equation (2), we obtain the following equations for the evolution of transmission and virulence:

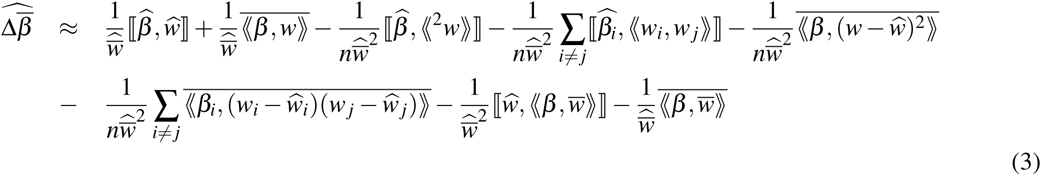

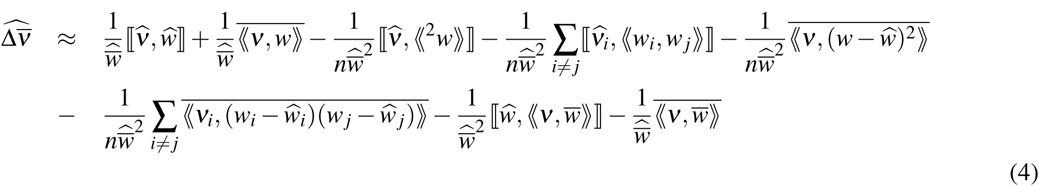

Note that absolute fitness, *w*, is a function of both transmission (*β*) and virulence (*ν*). We can write the linear approximation of *w* with respect to *β* and *ν* about origin as:

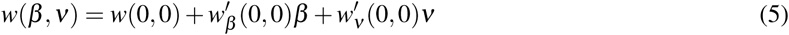

In the above equation, 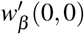 and 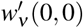 are the partial derivatives of *w* with respect to *β* and *ν*, respectively. For convenience, we use the notations 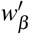; instead of 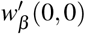 and 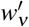 instead of 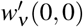. Substituting Equation (5) in to Equations (3) and (4), yields:

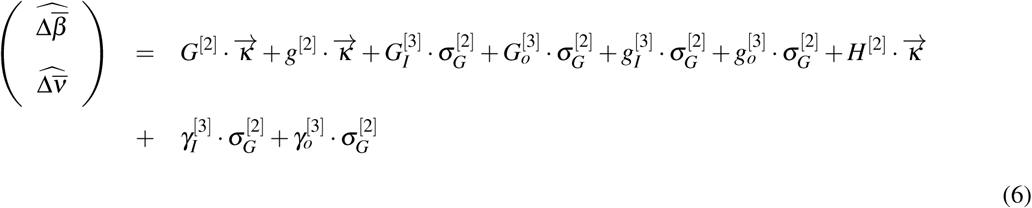

Detailed derivations are presented in Methods and Appendix 1. The terms on the righthand side of Equation (6) are written in terms of vectors, matrices, and tensors of degree 3. Figures (3,…,8) show schematic diagrams of tensors in the Equation (6). Below, we discuss the biological interpretations of each term of the Equations (3), (4), and (6).

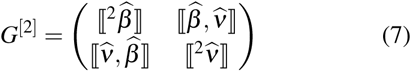

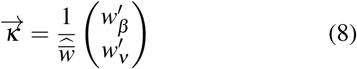

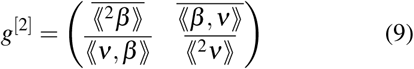

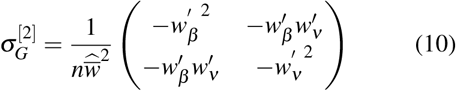

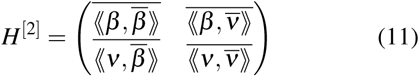

**Figure 3:**
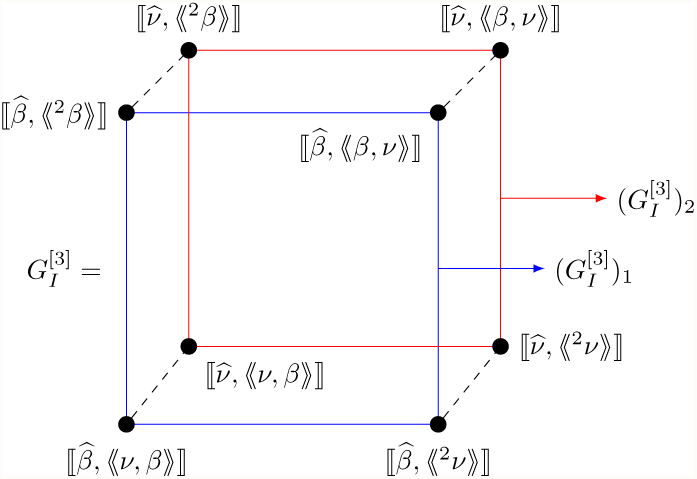
Schematic diagram of 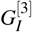

**Figure 4:**
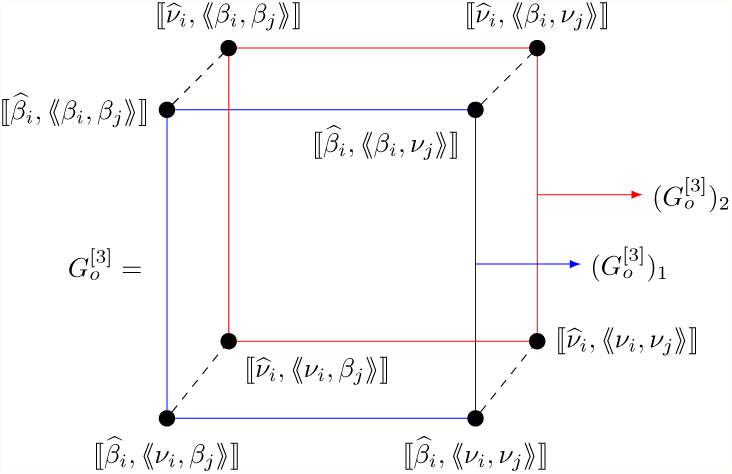
Schematic diagram of 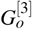

**Figure 5:**
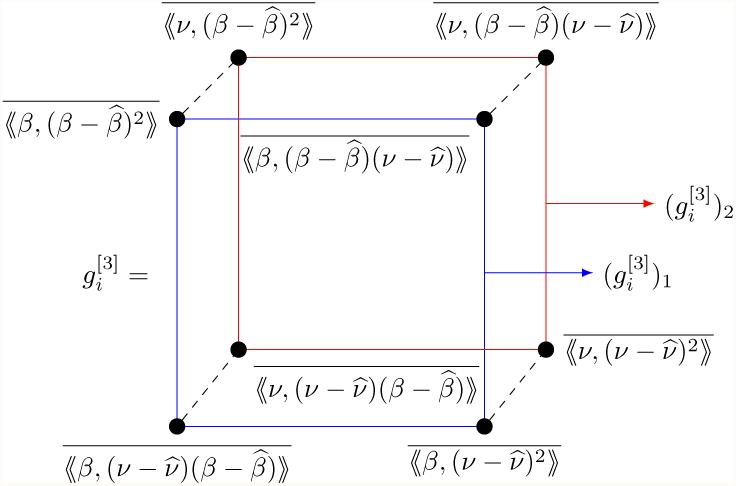
Schematic diagram of 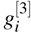

**Figure 6:**
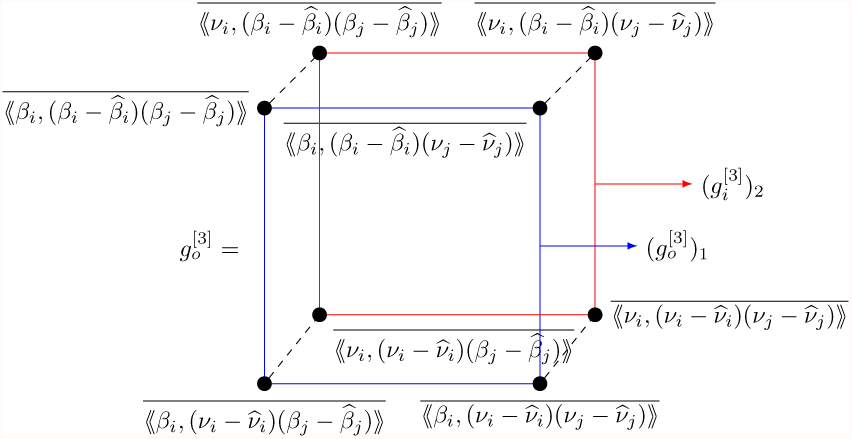
Schematic diagram of 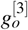

**Figure 7:**
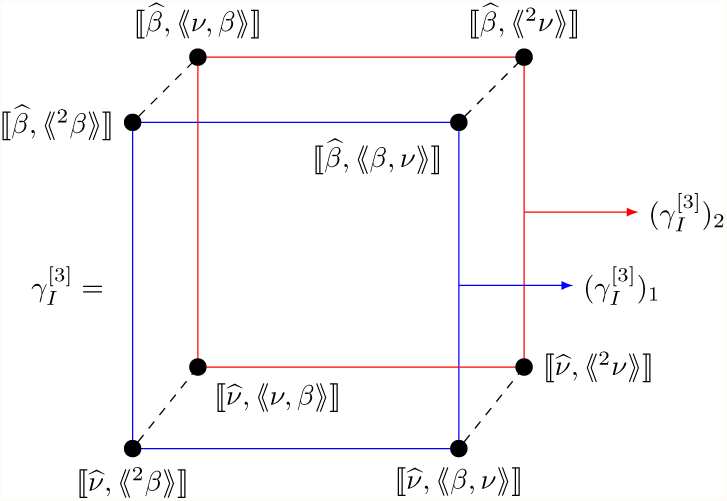
Schematic diagram of 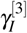

**Figure 8:**
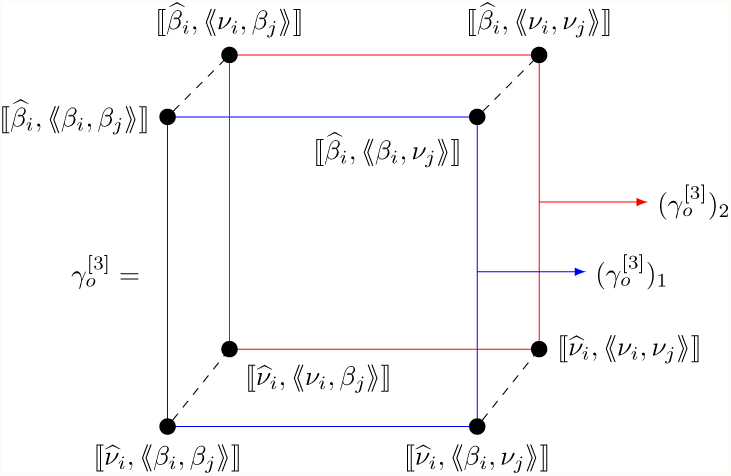
Schematic diagram of 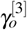

### Frequency Covariance versus Probability Covariance

The first terms on the right-hand side of the Equation (3) 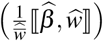 and Equation (4) 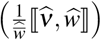 contain the frequency covariances between the expected absolute fitness of a pathogen strain and its expected transmission and virulence. This is the same as the covariance term in the original, deterministic, Price equation [18]. The first term on the right-hand side of the Equation (6) contains a matrix, *G*^[2]^, and a vector, 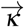, defined as:

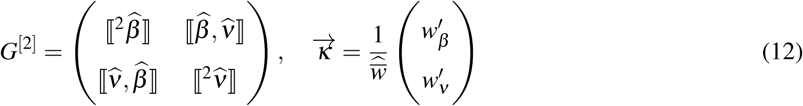

*G*^[2]^, the matrix of frequency covariances between our traits, is analogous to the standard *G* matrix in quantitative genetics [19]. It is multiplied by the fitness gradient, 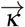, which shows the direction of maximum increase in fitness [19, 20]. The direction of evolution is influenced by both the fitness gradient, *κ*, and the frequency covariance matrix, *G*^[2]^, as shown in Figure (9). If 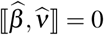, the directions of evolution is given by the selection gradient; increasing transmission and decreasing virulence. If 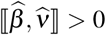, virulence may increase, despite the fact that it is selected to decrease. If 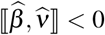, selection tends to increase the degree to which transmission increase and virulence decrease because the covariation between *β* and *ν* is aligned with the selection gradient action on them.

**Figure 9:**
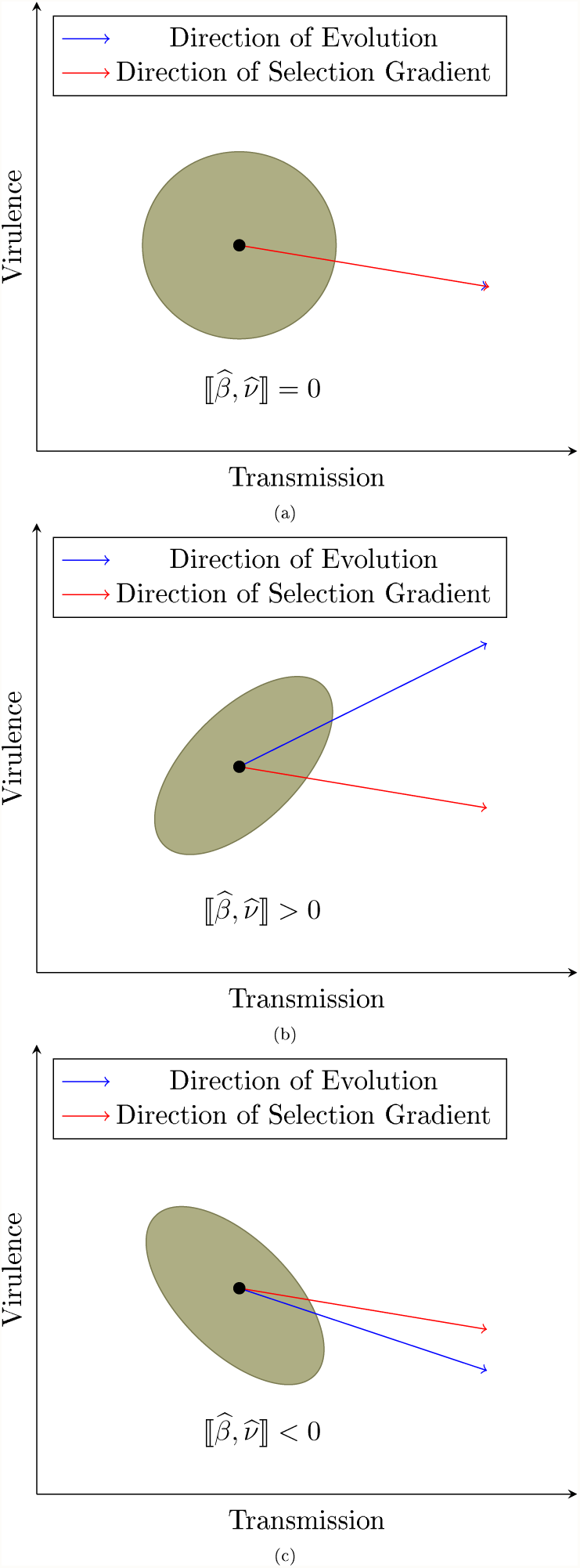
Covariance between traits can affect the direction of evolution. (a) Selection favors reducing virulence and since there is a zero frequency covariance between virulence and transmission, evolution points to the same direction as selection gradient. (b) There is a positive frequency covariance between virulence and transmission and selection tends to reduce the degree to which transmission can increase and virulence can decrease. (c): Frequency covariance between virulence and transmission is negative, and selection tends to increase the degree to which transmission can increase and virulence can decrease.

In the deterministic models such as [21, 22], the frequency covariance is the only term capturing the relationship between the fitness and phenotype. When phenotype and fitness are stochastic, however, two new kinds of terms appear, corresponding to evolutionary processes that are invisible to deterministic models.

When fitness and phenotype are random variables, they can covary for a single individual. This is captured by the second terms of the Equation (3), 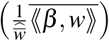 and Equation (4) 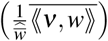. These terms contain the probability covariances between the absolute fitness of the pathogen strain with its transmission and virulence, averaged over the entire population.

The second term on the right-hand side of the Equation (6) also contains the fitness gradient, 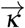, but here it is multiplied by a different matrix, *g*^[2]^, defined as:

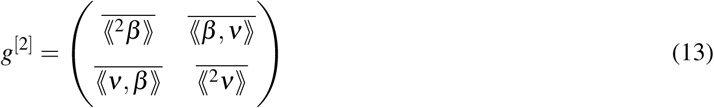

The matrix *g*^[2]^ contains the frequency means, across all pathogen strains, of these probability variances and covariances. The probability covariance between transmission and virulence can be positive or negative, depending on the biology of pathogen and host. For example, if the process of transmission requires harming the initial host, we would expect a positive probability covariance between virulence and transmission (Figure 10-A). Such is the case with amoebic dysentery, caused by the unicellular protozoan *Entamoeba histolytica*. In this case, the pathogen gets out of the host by inducing diarrhea [23], which can cause dehydration and be lethal if not treated. Increasing transmission of amoebic dysentery is thus likely to accompany increasing virulence [24]. By contrast, we expect a negative probability covariance between transmission and virulence in cases in which transmission of pathogens relies on a reasonably healthy host (Figure 10-B). Sexually transmitted diseases such as HIV infection should thus show a negative value of 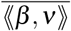.

**Figure 10:**
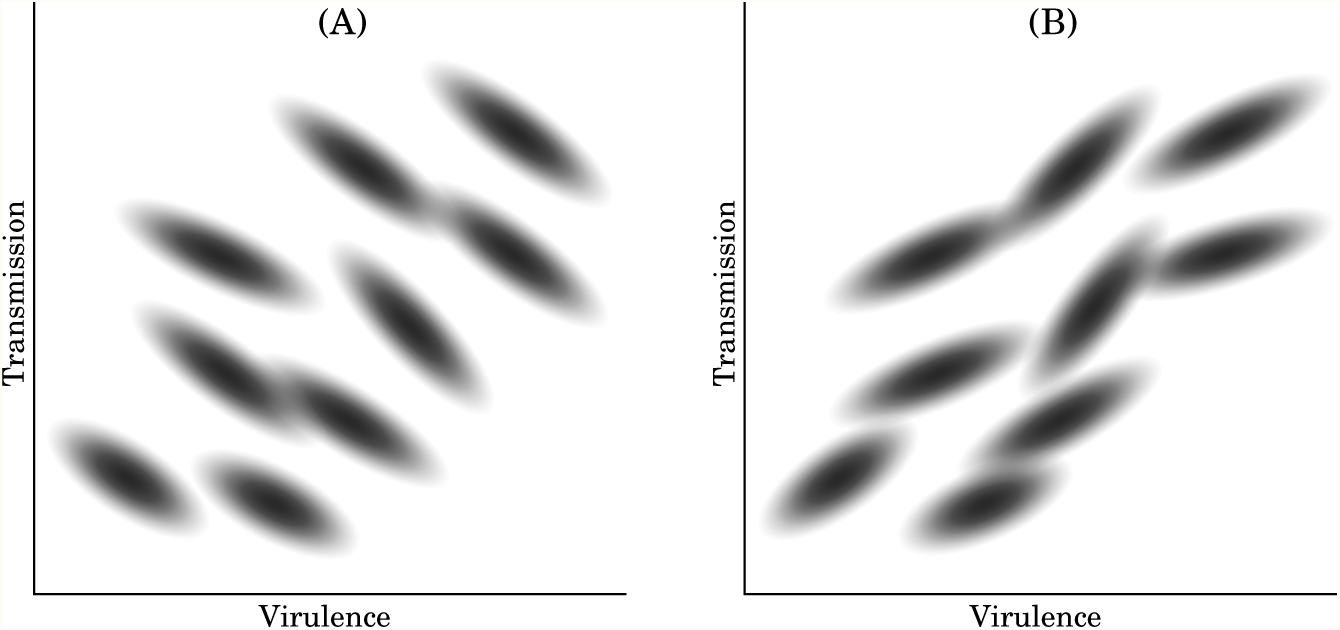
Two hypothetical cases of pathogen strains in which transmission and virulence are correlated. In Figure (A) we see a negative probability covariance between transmission and virulence that correspond to pathogen strains that are dependent on their hosts for transmission. But Figure (B) corresponds to pathogen strains with positive probability covariance between transmission and virulence. Those strains are not dependent on their hosts for transmission and generally the process of transmission includes harming hosts.

Equations (6) and (13) show that the expected change in mean transmission, 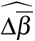, contains the term 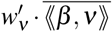. Similarly, the expected change in mean virulence, 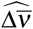, contains the term 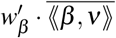. As we mentioned earlier, based on the biology of the pathogen 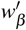 and 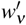 have different signs. For example, those pathogens that need reasonably healthy hosts for their process of transmission have 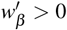 and 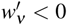. Thus a positive probability covariance between *β* and *ν* would inhibit the evolution of increased transmission and reduce the degree to which *ν* declines (possibly even causing it to increase). A negative value of 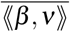 would have the opposite effect; amplifying both the effects of selection for increased transmission and reduced virulence.

To illustrate the difference between the frequency covariance, 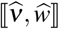, and the probability covariance, ⟪*ν, w* ⟫, Figure (11) shows a hypothetical case in which they have different signs. We consider a population of hosts that vary in immune resistance; with some having a strong immune response and some a weak response. Those pathogens that infect hosts with a strong response will tend to have both low expected fitness (*ŵ*) and low expected virulence 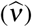, while those infecting hosts with weak immune response will have higher values of both *ŵ* and 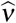 (black dots in the figure). We thus have a positive frequency covariance between expected fitness and expected virulence 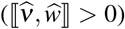. Considering any one pathogen within a particular host, however, the pathogen is likely to have higher fitness if it does not kill the host quickly. There is thus a negative probability covariance between fitness and virulence (⟪*w, ν* ⟫ < 0) for each individual pathogen.

The concept of a probability covariance between transmission and virulence is relevant to one of the central issues in the study of pathogen evolution: the idea that there are “tradeoffs” between transmission and virulence that constrain the evolution of both traits. Anderson and May assumed a positive association between transmission and virulence [25]. Subsequently, other authors have considered different tradeoff scenarios corresponding to different biological properties of pathogens and hosts [26, 27]. The kinds of tradeoffs described by these authors are more accurately captured by the probability covariance between *β* and *ν* ((⟪*β, ν* ⟫) than by the frequency covariance between their expected values. The frequency covariance 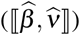 is influenced by the current distribution of pathogen strains and available hosts (Figure 11), whereas the probability covariance captures tradeoffs resulting from the basic biology of pathogen and host.

**Figure 11:**
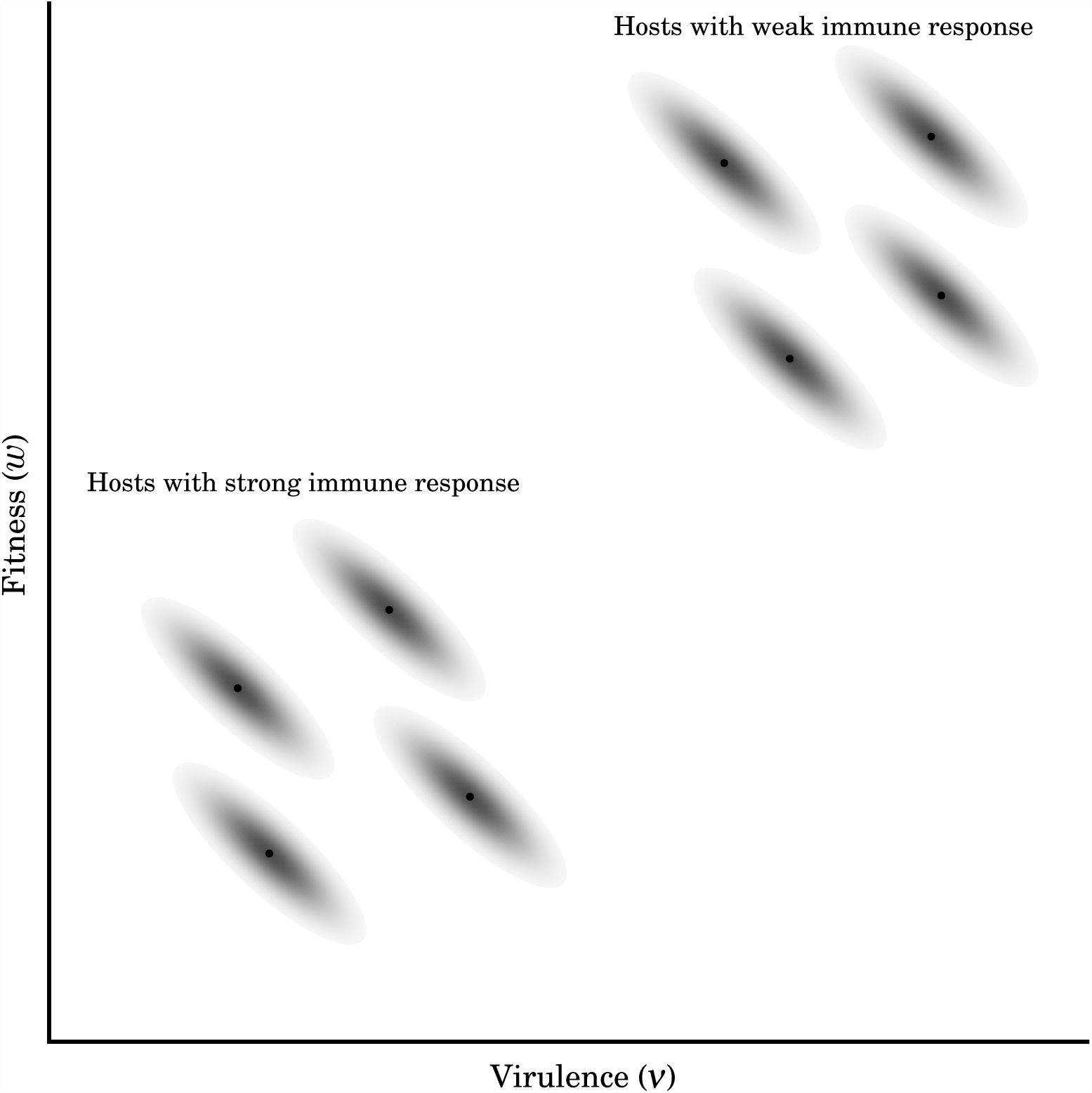
Probability and frequency capture different biological processes. A hypothetical case of eight different pathogen strains, in eight hosts, for which 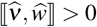, but ⟪*ν*_*i*_, *w*_*i*_ ⟫ < 0 for each strain (so 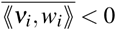). Each solid black dot represents the expected values of virulence 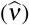 and fitness (*ŵ*) for that strain. Pathogen strains on the lower left corner show low expected virulence and low expected fitness since they happen to be in hosts with strong immune responses. The strains on the upper right corner of the figure are in hosts with weak immune systems, leading to high expected virulence but also higher fitness of the pathogen. Within any one host, the pathogen has a greater chance of transmission if it does minimal harm to its host; leading to the negative probability covariance between fitness and virulence for each strain.

Measuring the probability covariance between *β* and *ν* would, ideally, involve using replicate clones of both pathogen and host, to minimize variation due to genetic differences. We know of no experimental studies have specifically done this. However, some studies using the *Daphnia magna*-*Pasturia ramosa* system have provided data from which 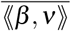 can be estimated [13, 14]. In one study, Vale et. al estimated the covariance between transmission potential and virulence for bacteria in clones of *Daphnia magna* (Figure 12) [13]. (To increase the number of datapoints, these authors infected the clones of *Daphnia magna* by different strains of *Pasturia ramosa*, so this is not a pure estimate of probability covariance.) One interesting conclusion from these studies is that the value of ⟪*β, ν* ⟫ can change sign as a result of environmental changes. Specifically, the covariance between *β* and *ν* was negative when hosts were given ample food resources, but became positive when hosts were severely nutrient stressed [13]. Thus, physiological stress on the host can lead to the evolution of increased virulence of the pathogen.

**Figure 12:**
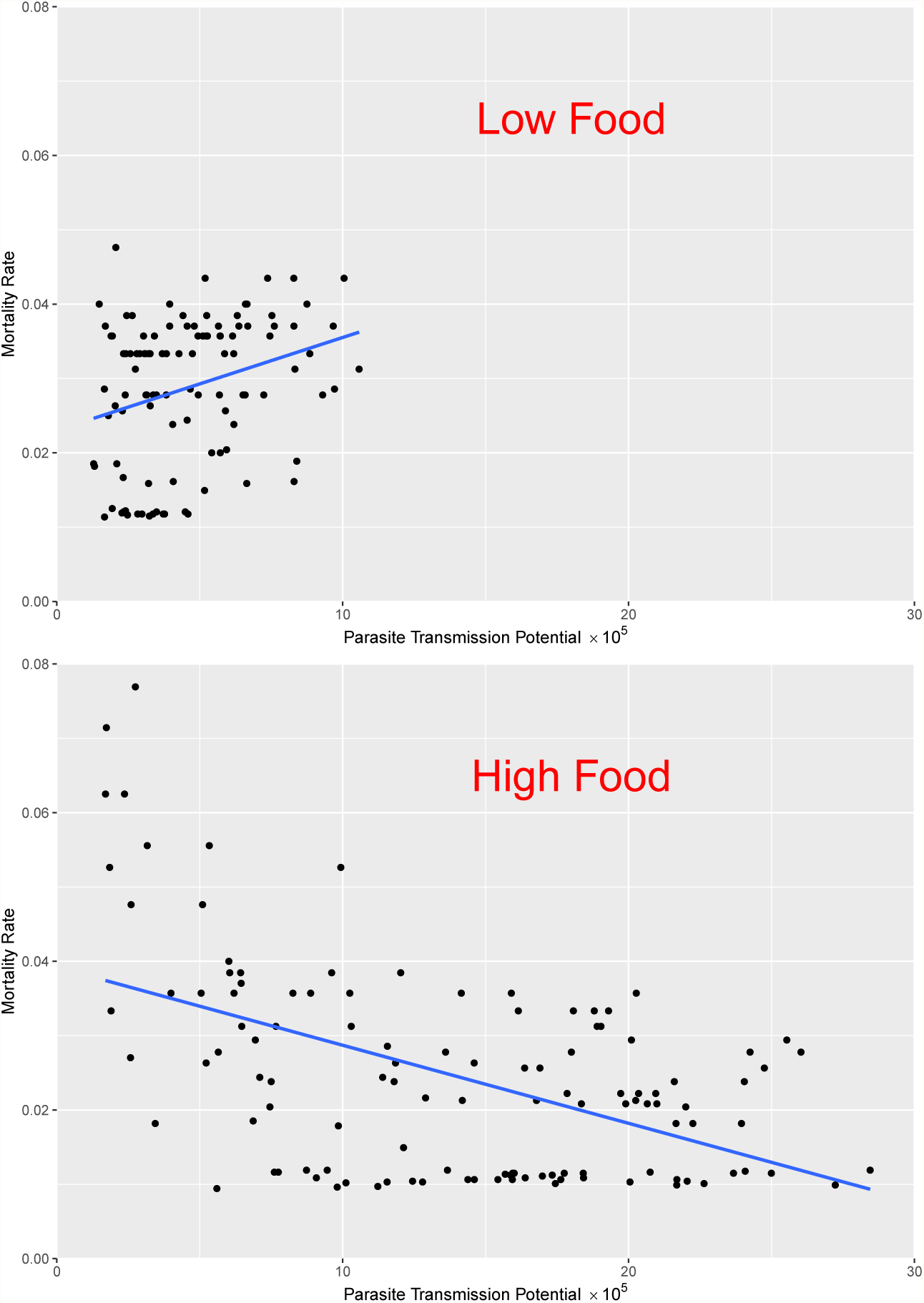
The relation between *Daphnia magna* mortality rate and *Pasturia ramosa* transmission potential. The top figure shows that when clones of the hosts are severely nutrient stressed, the covariance sign becomes positive. But when hosts are given ample food resources, the sign changes and becomes negative. One striking conclusion from this study is that the value of ⟪ *β, ν* ⟫ can change sign not only as a result of the biology of pathogen, but also as a result of environmental changes. The blue line shows the least square regression line. Figure is based on data by [13].

### Directional Stochastic Effects

The terms discussed above result from probability covariation between fitness and descendant phenotype. The second class of evolutionary phenomena that appears when we make fitness stochastic – directional stochastic effects – result from covariation between individual fitness (*w*) and mean population fitness 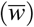 (written as 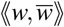 in the third term on the righthand side of Equation (2)).

Figure (13A) illustrates the basic mechanism of directional stochastic effects. If all else is held equal, the magnitude of evolutionary change is inversely proportional to mean population fitness. This is a general property of evolutionary models, manifest in the fact that population genetic models for change in allele frequency, and quantitative genetic models for change in mean phenotype, always contain a 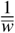 term (unless it is assumed that 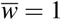). Figure (13A) shows a hypothetical case for a population of two individuals that have the same expected fitness, but different variances. In the example, there is an equal probability that mean phenotype will increase or decrease. when 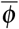 increases, though, the absolute magnitude of change is greater than when it decreases (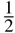 *vs.* 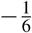). The reason for this discrepancy is that 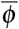 increases when 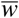 is low, and decreases when 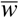 is large.

**Figure 13:**
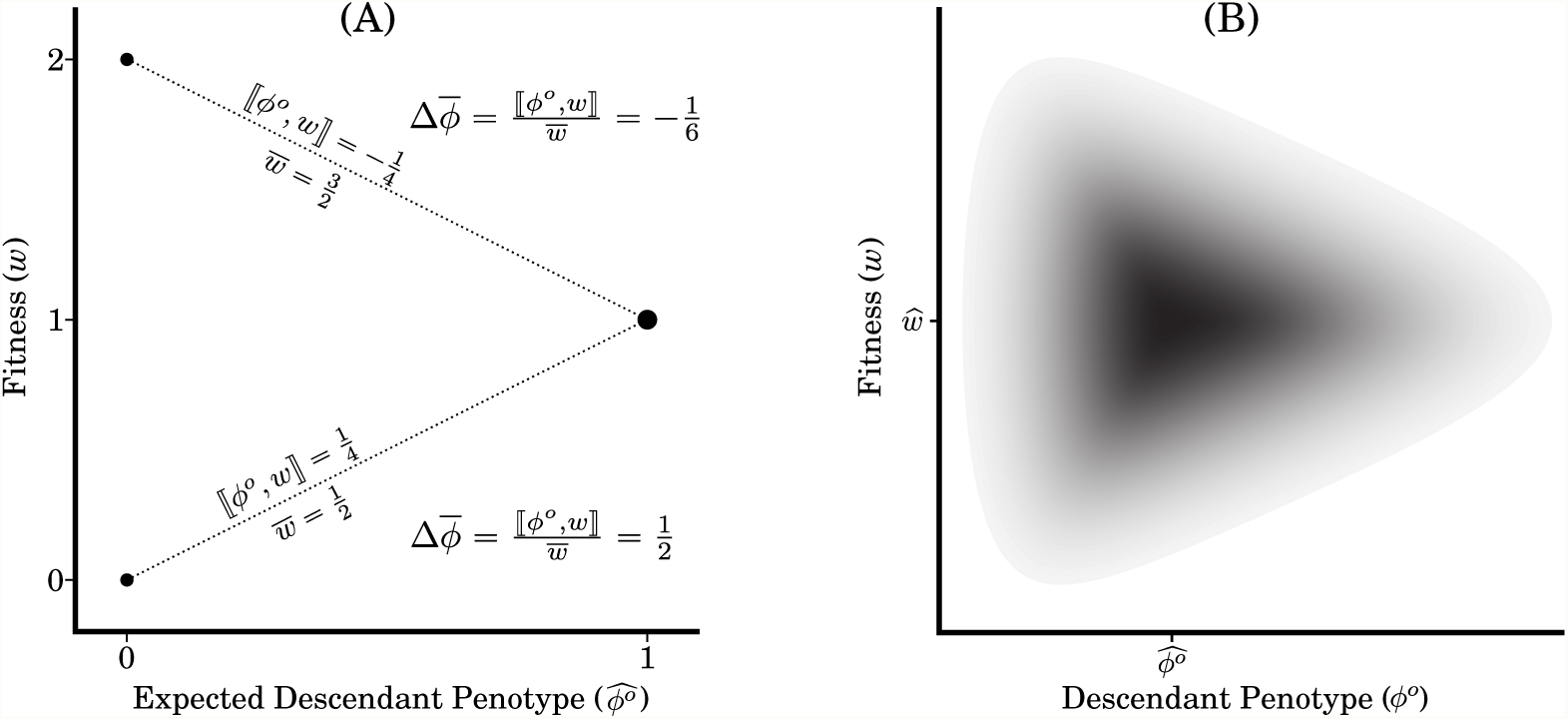
Directional Stochastic Effects. (A) A population of two sets of individuals with different offspring phenotypic values, 0 and 1, havin <u>g</u> different fitness distributions. When those with 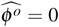 have two offspring each, 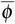 declines by 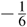, when they have zero offspring each, 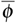 increases by 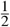; for a total expected change of 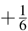. Though the strength of selection (captured by **⟦** *ϕ*^*o*^, *w* **⟧**) is the same, and in opposite directions, the change in 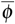 is different in the two cases because 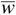 is different. (B) Joint fitness and offspring phenotype distribution for an individual in which **⟪** *ϕ*^*o*^, (*w* − ŵ)^2^**⟫**) is nonzero. This probability distribution of *w* and *ϕ*^*o*^ within an individual leads to the same evolutionary effect as does the frequency distribution across individuals shown in (A).

Because any individual’s fitness is a component of mean population fitness, the probability covariance between *w* and 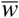 can be broken up into two components; one involving the variance in an individual’s fitness, and the other involving the probability covariance between that individual and others in the population. For the *i*^*th*^ pathogen strain, we can write:

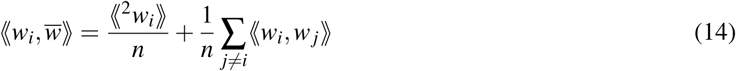

This is the origin of the 3^*rd*^ and 4^*th*^ terms in Equations (3) and (4).

The third terms in the Equations (3) and (4), 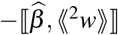 and 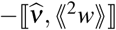, are negative, meaning that they act to pull the population towards phenotypes that have minimum variance in fitness (⟪^2^*w* ⟫). Thus, if 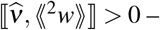 meaning that expected virulence covaries positively with variance in fitness – then this directional stochastic effect will tend to reduce the expected change in virulence. Note that this effect is distinct from selection proper, which is captured by the 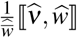 term. As a result, it is possible for a particular strain of the pathogen to have a lower expected fitness, but to nonetheless be expected to increase, if it also has a much lower variance in fitness than do other strains. The magnitude of directional stochastic effects is greatest when either the number of pathogen strains (*n*) or the expected pathogen growth rate 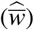 is small. We thus expect these effects to be most important at times of low pathogen diversity – such as when a pathogen is introduced into a new host population – or when the pathogen population within a host is declining – such as when it is under attack by the host’s immune system. (Note that the directional stochastic effects are different from drift. Drift is nondirectional, in the sense that the expected frequency change of the phenotype due to the drift alone is zero).

Intuitively, the reason that there is a directional stochastic effect acting to reduce variance in fitness is that a strain with high variance (high ⟪^2^*w* ⟫contributes disproportionately to the variance in mean population fitness, 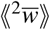. As a result, when that strain with a high value of *ϕ* and high ⟪^2^*w* ⟫ happens to have a particularly high fitness, 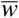 also tends to be high – reducing the magnitude of increase. In contrast, when that strain has a lower value of *w*, then 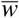 also tends to be low – leading to a large decline in 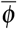. This is the case for the individual with *ϕ* = 0 in Figure (13A).

The third term on the right-hand side of the Equation (6), 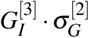, is the inner product of a third order tensor, 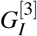, and a second order tensor, the matrix 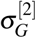. These can be written as follows (where 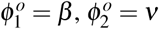,):

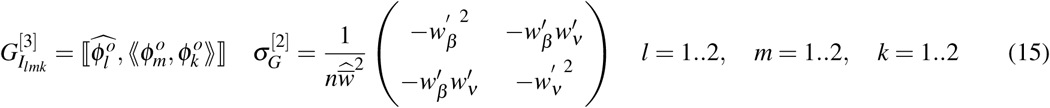

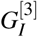 is a three dimensional array, with element (*l, m, k*) defined by Equation (15) and its schematic diagram is represented in Figure (3). The term 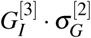 captures directional stochastic effects, as discussed earlier. The elements of 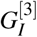 are components of the 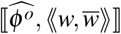 term from Equation (2). Taken together, the combined elements of 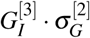 push the population towards phenotypes that minimize the variance in individual fitness and the covariance between the fitness values of different individuals.

To demonstrate how expected virulence can be correlated with probability variance in pathogen strains (**⟦** *ν*, ⟪^2^*β* ⟫ **⟧**), we can use the hypothetical example based on well known example of Myxoma virus [28]. Experimental data show that when Myxoma virus first introduced in 1950 to control the number of invasive rabbits in Australia, the early generations of the pathogen were highly virulent, killing about 99% of their hosts [28]. Later generations of pathogen, however, evolved to be less virulent. The possible explanation of this is to be found in the mechanism of pathogen transmission. When there is an ample number of hosts in the population, the virulent strains of the pathogen are favored by selection because they grow faster inside their host which in turn can boost their transmission. But as the number of the hosts declines dramatically, the contacts between hosts also decline, and the virulent strains no longer have an advantage since most of the hosts die before contacting with another host.

Now consider a small population of rabbits such that there is a high variation in contacts between them that sometimes hosts form groups. Introduce two different strains of Myxoma virus, with high and low virulence to this population. As a consequence of growing faster inside its host, the high virulent strain can benefit a higher transmission from a current host to a new host only if the current host stays alive long enough for transmission. Therefore, a highly virulent pathogen can sometime gain high transmission and sometimes no transmission. But low virulent pathogen grows lower inside its host and therefore it has no chance of very high transmission. On the other hand, since it imposes low pathology on its host, a low virulence strain usually has some moderate level of transmission. In this situation we have a positive correlation between virulence and probability variance of transmission. As a result, directional stochastic effects, tend to pull the population toward low virulence strains that have minimum variance in transmission.

Equation (14) shows that there is another way in which an individual’s fitness can covary with mean fitness – if it covaries with the fitness of others. This is the source of the fourth terms in Equations (3) and (4) (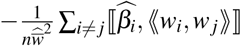 and 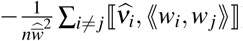). These terms capture the effects of covariance between the fitness of different pathogen strains on the evolution of transmission and virulence. An example of a case in which we might expect 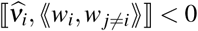) is a case in which there are multiple strains competing within the host, such that the most virulent strain is the best competitor (so that other strains tend to do poorly when it does well). This would make the fourth term on the righthand side of Equation (4) positive; contributing to an increase in mean virulence.

The fourth term on the right-hand side of the Equation (6), 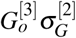, like the third term, is defined in terms of tensor notation: (where 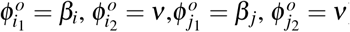):

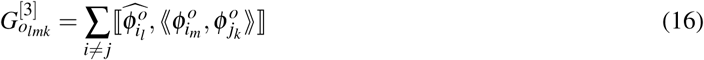

Figure (4) shows that the components of this tensor involve the interactions between the phenotypes (*β* and *ν*) of the *i*^th^ pathogen strain and every other strains. The biological interpretation is similar as what described for Equation (14).

The third and fourth terms in Equations (3) and (4), discussed above, appear whenever fitness is a random variable. If descendant phenotype is also a random variable, as we are assuming here, then we also encounter terms containing the probability covariance between phenotype and mean population fitness 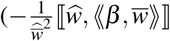 and 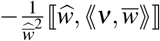 in our example) which are the seventh terms in the Equations (3) and (4). The interpretation of these is similar to that of the cases containing 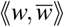, except here it is a probability covariance between phenotype and mean fitness that influences the magnitude of change when it is associated with the expected fitness of individuals. The eighth and ninth terms of the Equation (6), 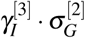 and 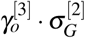, are in terms of *β* and *ν* and correspond to the effects of the seventh terms of the Equations (3) and (4)(where 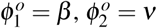):

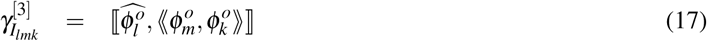

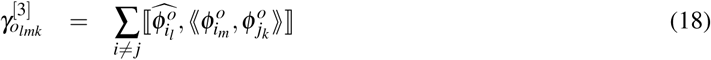

Figures (7) and (8) show a schematic diagram of the tensor 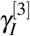 and 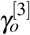. It is interesting to note that some terms such as 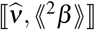 that in previous terms had direct roles in the evolution of *ν*, now have direct influence on the evolution of *β*.

The fifth terms of the Equations (3) and (4), 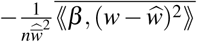 and 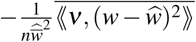 involve an association between a stochastic phenotype and uncertainty in fitness. (They are thus the stochastic phenotype analogues of the third terms). Figure (13B) shows a case in which ⟪ *ϕ*^0^,(*w* − ŵ)^2^⟫is nonzero. Note that the expected value of (*w* − ŵ)^2^ is just the probability variance in fitness. Saying that ⟪ *β*,(*w* − ŵ)^2^⟫ < 0is equivalent to saying that, for a particular pathogen strain: when that strain has higher than expected transmission (*β*), it also has lower uncertainty in fitness. The fifth and the sixth terms of the Equation (6) correspond to the effect of this term.

The fifth of the Equation (6), 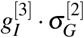, is made up of the following terms (where 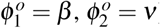):

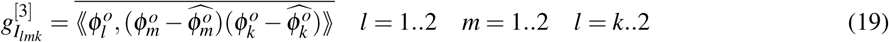

and the Figure (5) corresponds to the tensor 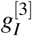:

The sixth term of the Equation (6), 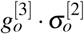, is made up of the following terms (where 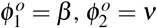):

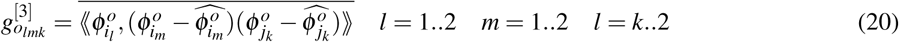

and the Figure (6) corresponds to the tensor 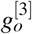:

A similar interpretation applies to the terms 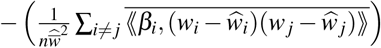 and 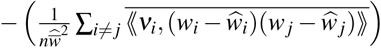. Here, it is an association between stochastic phenotype and the similarity in fitness between different individuals (or strains) that influences evolution.

Finally, the terms 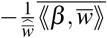 and 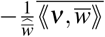 capture the direct correlation between stochastic phenotype and mean population fitness, averaged over the entire population. As in the examples above, these terms are negative because high values of 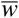 reduce the magnitude of evolutionary change, so the magnitude of change in a trait is reduced when that trait covaries with 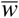.

The last term of the Equation (6) capture the direct correlation between the individual stochastic phenotype and population mean phenotype:

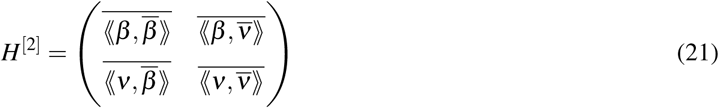

Equation (6) is based on a general linear relationship between transmission, virulence, and fitness (Equation (5)). If we know, for a particular pathogen, how fitness is related to *β* and *ν*, we can substitute that function into Equations (3) and (4) to derive a specific model for that pathogen. In the next section, we do this for the widely studied SIR model of pathogen dynamics.

### Special Case: The SIR Model

The SIR model is a compartmental epedemiological model in which we follow the dynamics of three different categories of hosts: those that are uninfected but susceptible to the pathogen (*S*), those that are infected (*I*), and those that have recovered from infection (*R*). For pathogen strain *i*, the continuous time *SIR* model is:

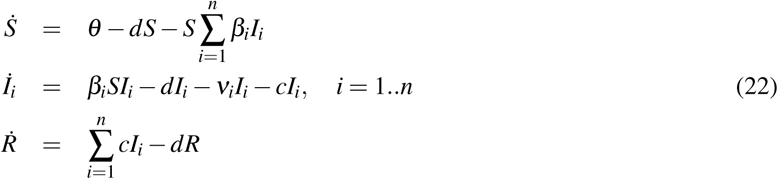

In Equations (22), *I*_*i*_ represents the density of individuals infected by the i*th* strain of the pathogen, *β* is transmission rate, *ν* is virulence (the degree to which the infection increases the death rate of infected hosts), *d* is the background death rate (in the absence of infection), and *c* is the “clearance” rate (the rate at which infected hosts eliminate the infection, such as through immune response). A dot over a variable indicates a time derivative. (See Table 1.)

Day and Gandon developed a way to apply the deterministic Price equation to the evolution of pathogens that follow *SIR* dynamics [21, 22]. They assume that a host can harbor, at most, a single pathogen strain at any given time. In this case, the fitness of pathogen strain *i* is equal to the per capita growth rate of infected hosts, *I*_*i*_, that carry strain *i*. Using Equations (22), we can write the fitness of pathogen strain *i* as a function of transmission (*β*) and virulence (*ν*):

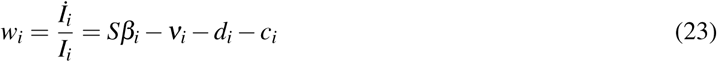

Note that using a different epidemiological model would give us a different equation for the absolute fitness.

We consider a pathogen that has just infected a host, and treat descendant phenotype (*ϕ*^*o*^ in the general equation) as the value of *β* or *ν* when that pathogen’s descendants are transmitted to another host. We thus essentially treat *β* and *ν* as having heritabilities of 1. By contrast, *d* and *c* are properties of the host, and are thus environmental variables from the perspective of the pathogen. We will include them because they may interact with *β* and *ν* to influence pathogen evolution, but we assume that *d* and *c* are not heritable by the pathogen.

We introduce stochasticity by treating the parameters *β, ν, d*, and *c* in Equation (23) as random variables. For simplicity, and consistent with the assumption that a host can harbor only one strain at a time, we will assume that different strains are stochastically independent of one another - meaning that the fourth and sixth terms of Equations (3) and (4) are zero.

Substituting Equation (23) into Equations (3) and (4) and rearranging the terms extensively, yields the following equation for the vector of expected changes in mean transmission, and virulence, natural mortality, and clearance rate over one generation (see Appendix 2):

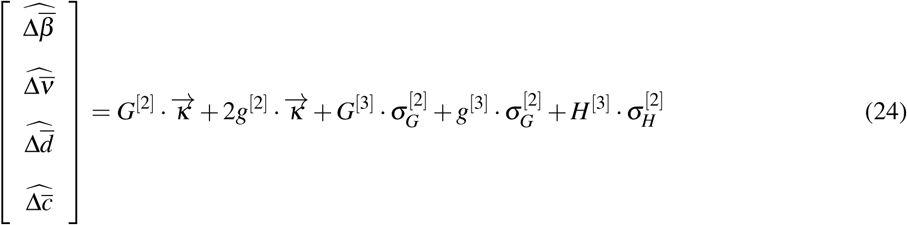

The terms on the righthand side of Equation (24) are written in terms of vectors, matrices, and tensors of degree 3. Below, we discuss the biological meaning of each term in turn.

The first term on the right-hand side of the Equation (24) contains a matrix, *G*^[2]^, and a vector, 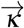, defined as:

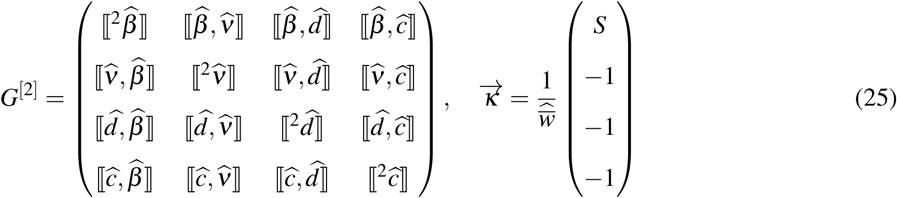

*G*^[2]^, the matrix of frequency covariances between our traits, is analogous to the standard *G* matrix in quantitative genetics. It is multiplied by the fitness gradient, 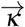, which shows the direction of maximum increase in fitness [20, 19]. The matrix *G*^[2]^ will appear in any model with parameters *β, ν, d*, and *c*. The specific form of 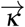 given in Equation (25), however, is specific to the SIR model (Equations 22) – a different model would yield a different 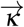.

For the case of the *SIR* model, the form of 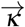 shown in Equation (25) shows that selection always favors increasing the average transmission with a strength of 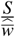 (where *S* is the density of uninfected but susceptible hosts), while decreasing the average virulence, background mortality, and recover rate, with a strength of 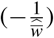. How this selection influences evolution is of course also influenced by *G*^[2]^. A positive covariance between *β* and *ν* will tend to reduce the degree to which transmission can increase and the degree which virulence can decrease.

Note that the expected change in mean transmission, 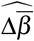, contains the terms 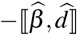 and 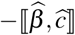. This illustrates the importance of including the “environmental” factors *d* and *c* in our analysis.

In the absence of mutation, 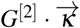 is the only term that would appear in a deterministic model, and is equivalent to the selection term in the models of [21, 22]. All of the other terms in Equation (24) contain variances and covariances of random variables; and are thus invisible to deterministic models.

The second term on the right-hand side of the Equation (24) also contains the fitness gradient, 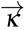, but here it is multiplied by a different matrix, *g*^[2]^, defined as:

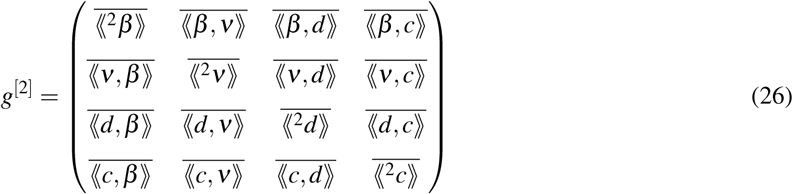

Since *β, ν, d*, and *c* are random variables, each has distribution of possible values for any one pathogen strain; they can therefore covary for a single pathogen. The matrix *g*^[2]^ contains the frequency means, across all pathogen strains, of these probability variances and covariances.

Equations (24) and (26) show that the expected change in mean transmission, 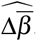, contains the term 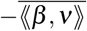. The expected change in mean virulence, 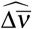, contains the term 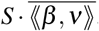. Thus a positive probability covariance between *β* and *ν* would inhibit the evolution of increased transmission and amplify the evolution of increased virulence. A negative value of 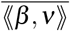 would have the opposite effect; amplifying the effects of selection for increased transmission while reducing the evolution of increased virulence.

The third term on the right-hand side of the Equation (24), 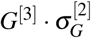, is the inner product of a third order tensor, *G*^[3]^, and a second order tensor, the matrix 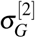. These can be written as follows (where 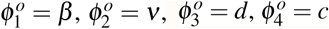):

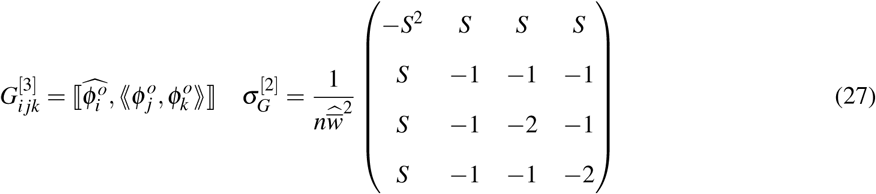

*G*^[3]^ is a three dimensional array, with element (*i, j, k*) defined by Equation (27) (Figure 14). The term 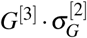 captures directional stochastic effects, as discussed in the previous section. The elements of *G*^[3]^ are components of the 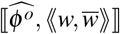 term from Equation (2). Taken together, the combined elements of 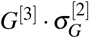 push the population towards phenotypes that minimize the variance in individual fitness and the covariance between the fitness values of different individuals.

**Figure 14:**
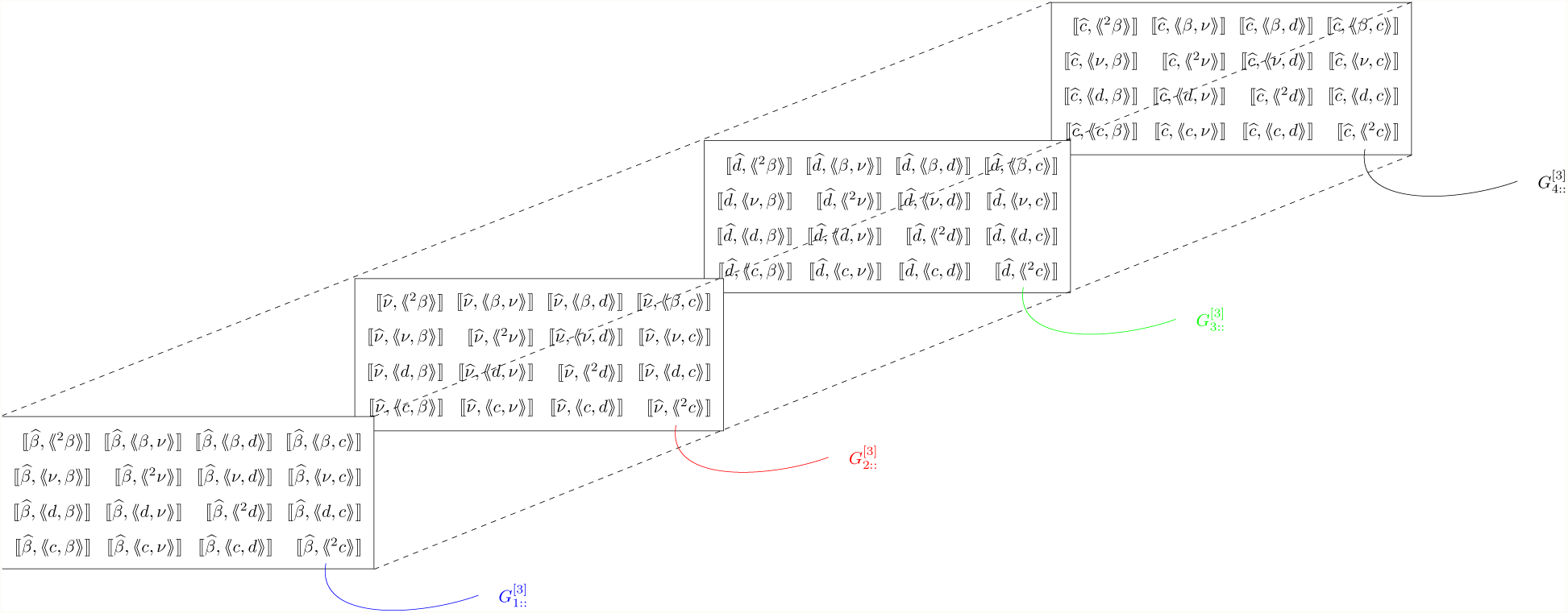
An illustration of third order tensor *G*^[3]^

Note that the only positive terms in 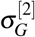 are terms that multiply elements of *G*^[3]^ that contain ⟪ *β, ν* ⟫, ⟪ *β, d* ⟫, or ⟪ *β, c* ⟫. This means that either *β* or *ν* will tend to increase if it has a positive frequency covariance with ⟪ *β, ν* ⟫, ⟪ *β, d* ⟫, or ⟪ *β, c* ⟫. The reason is that *ν, d*, and *c* all contribute negatively to fitness while *β* contributes positively.

The fourth and the fifth terms of the Equation (24), 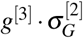 and 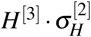, are made up of the following terms:

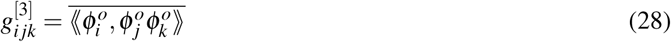

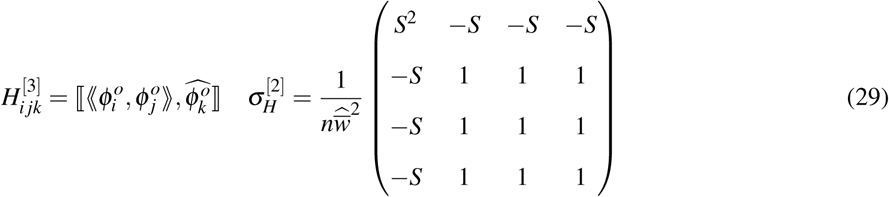

Though complicated, these terms all derive from the covariance between individual phenotypes and mean population fitness.

## Discussion

Uncertainty in both fitness and offspring phenotype can influence pathogen evolution in a variety of ways. This is clear from the fact that in the general equations for evolution of transmission and virulence, Equations (3) and (4), all of the terms after the first one, in each equation, contain probability operations and would thus not appear in a deterministic model. Though the equations contain many stochastic terms, the stochastic components of directional evolution discussed in this chapter all fall into two general categories: Probability covariances between traits, and directional stochastic effects.

Probability covariances between traits arise whenever the traits that influence fitness are random variables – meaning that they have a distribution of possible values. This is definitely the case for transmission and virulence of a pathogen since, when a pathogen infects a host, we can not say with certainty how long the host will survive or how many others it will infect.

The concept of a probability covariance between transmission and virulence is relevant to one of the central issues in the study of pathogen evolution which is the idea that there are “tradeoffs” between transmission and virulence that constrain the evolution of both traits. We argue that the kinds of tradeoffs between the phenotypes of the pathogen are more accurately captured by the probability covariance between them than by the frequency covariance between their expected values. The frequency covariance is strongly influenced by the current distribution of pathogen strains and available hosts, but the probability covariance captures tradeoffs resulting from the basic biology of pathogen and host. Experimental evidence suggests that the sign of the probability covariance depends not only on the biology of the pathogen, but also on the physiological state of the host.

The second way in which stochasticity can influence directional evolution is through directional stochastic effects. These result from the fact that the same strength of selection (the relation between expected fitness and phenotype, measured either as a covariance or a regression) will produce different magnitudes of evolutionary change depending on the mean population fitness 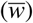, which is itself a random variable. The third through eighth terms in Equations (3) and (4) are all related to this kind of evolutionary process.

The simplest directional stochastic effect favors phenotypes that have relatively low variance in fitness (corresponding to the 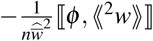 term in Equations (3) and (4); (note that this is not the same as having high geometric mean fitness [2]). Pathogen fitness involves both evading the host’s immune system – which is likely to exhibit stochastic variation both within and between hosts [29, 30] – and transmission from one host to another, which is likely to add further stochasticity. It is thus likely to be very unpredictable. A trait that reduces this uncertainty, even at the expense of reducing expected fitness, could thus spread – especially when the number of pathogen strains is small (small *n* in Equations (3) and (4)) or the population of pathogens is declining (small 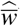).

Another way in which a trait can increase due to directional stochastic effects is through a negative probability covariance between the fitness of individuals possessing it and other individuals in the population. (Terms containing ⟦ *ϕ*_*i*_, (*w*_*i*_, *w*_*j*_) ⟧ in Equations (3) and (4)). For pathogens, this effect would favor a strain that deals differently with the host’s immune system than do other strains, even if it does no better on average. This effect would thus be a diversifying force with regard to how pathogens interact with their hosts.

Note that this approach to introducing stochasticity is different from that used in stochastic differential equations (SDE). In SDE, one makes the main variable stochastic by assigning a variation to it using a Wiener process, while other variables are kept deterministic [31]. For example, a SDE model for the evolution of virulence might treat transmission and overall fitness as fixed values, while assigning random normally distributed noise to *ν*. By contrast, our approach treats virulence, transmission, and fitness as random variables, which can have any distributions of values and can covary with one another.

Making our evolutionary models stochastic by allowing traits such as transmission and virulence to be random variables substantially increases the complexity of our results. Upon examination, however, the new terms that appear all correspond to real biological phenomena that would be missed by a purely deterministic analysis.

## Methods

The relative fitness of an individual, 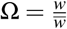 conditional on 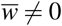, is a ratio of correlated random variables. In order to evaluate its expected value or its probability covariance with other random variables, we need to expand it in such a way that all random variables are in the numerator. Following [17], we define 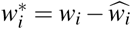, and expand Ω_*i*_ as:

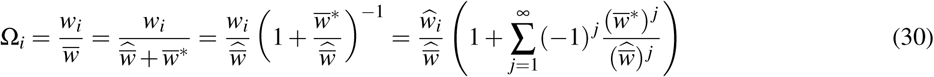

We now find the expected value, 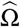 as [17]:

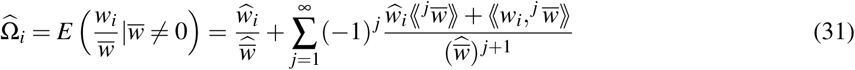

For a population of size *N*, the *j*^*th*^ order terms in these series tend to scale as 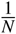, so we approximate 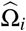by setting *j* = 1. This yields:

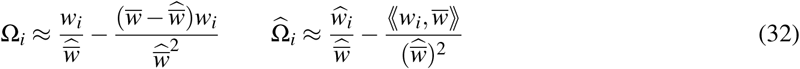

Using Equation (32), we write 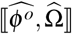 from Equation (1) as follow:

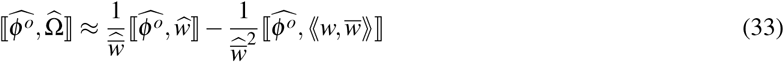

we also use Equation (32) and obtain:

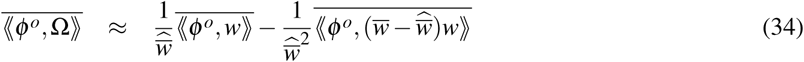

The second term of the Equation (34) can be manipulated as follow:

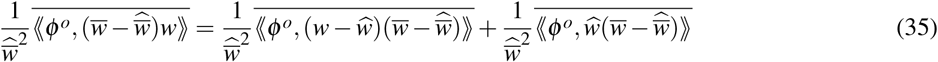

Recall the formula 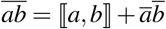 which bar indicates the average over all individuals in population. Therefore, the second term on the right-hand side of the Equation (35), can be expanded as follows:

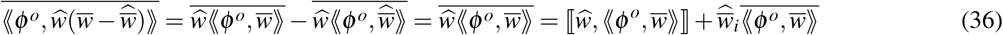

Therefore, we obtain:

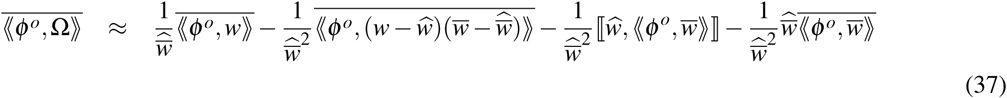

and substituting Equations (33) and (37) in Equation (1) yields Equation (2).

The third and fourth terms of the Equation (2) can further expanded as:

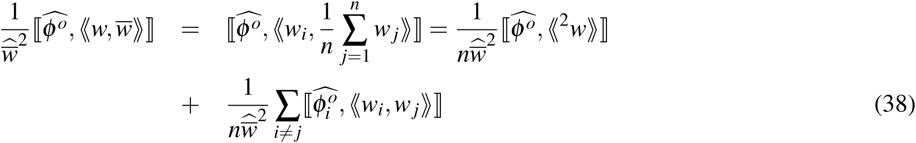

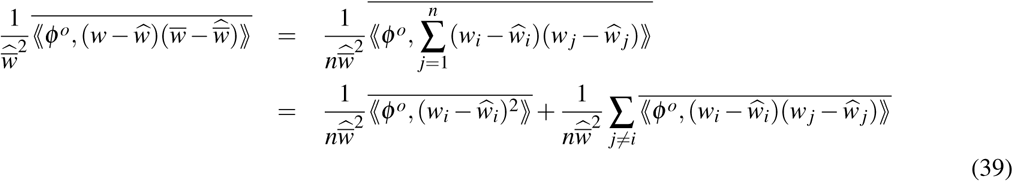

and substituting these equations in Equation (2) and *β* and *ν* for offspring phenotype, results Equations (3) and (4).

From Equation (5) we derive the equation for fitness of pathogen strain *i* as:

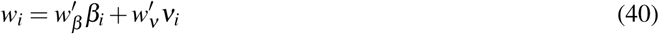

We present Equation (6) with the tensor notations. For the strain *i*, we define the random variables 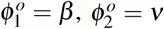. The covariance terms on the right-hand sides of Equations (6) can be grouped into 2 × 2 degree 2 and 2 × 2 × 2 degree 3 tensors (note that a tensor of degree 2 is just a matrix). The elements of each tensor can be shown as follow:

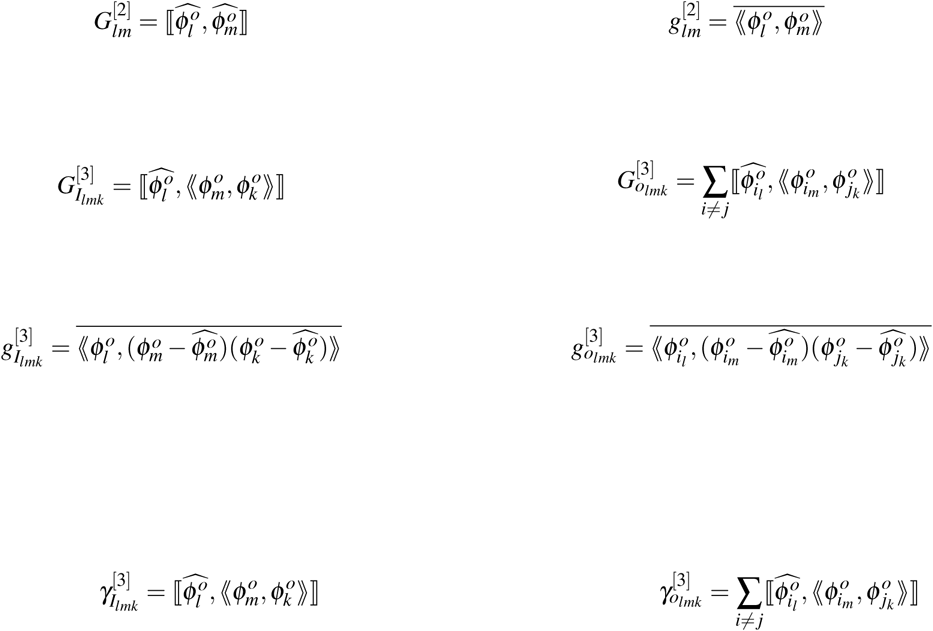

where *i* = 1…*n, j* = 1…*n, l* = 1…2, m=1…2, *k* = 1 2. Also we obtain:

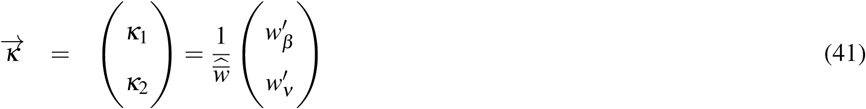

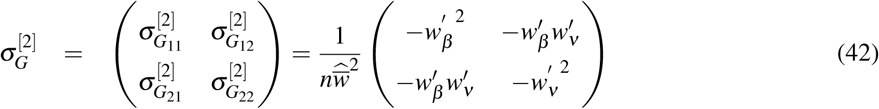

The tensor inner products on the right-hand side of equation (6) is 2 by 1 vector and can be written as follows:

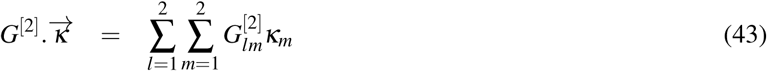

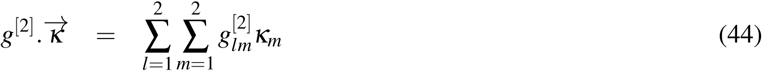

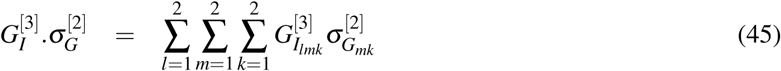

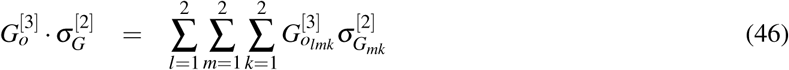

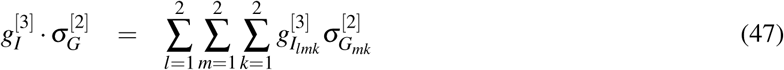

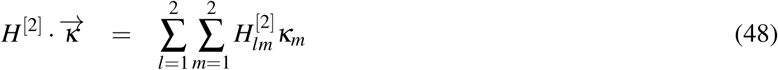

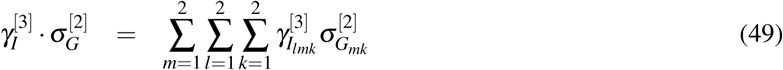

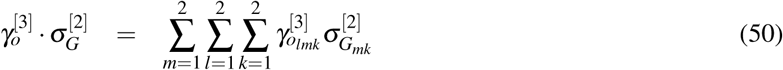

## Appendix1

We will use the following identities in manipulating frequency and probability operations:

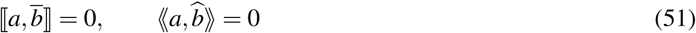

The following theorem is useful for manipulating covariance of products:

### Theorem 1.

i. *For three arbitrary random variables a, b, and c we have the following identity*

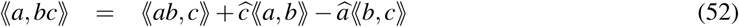
ii. *If c is independent from a, b, and ab, Equation (52) collapses to:*

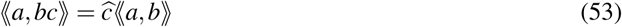

*Proof.*

i. We extract the right side of the Equation (52)

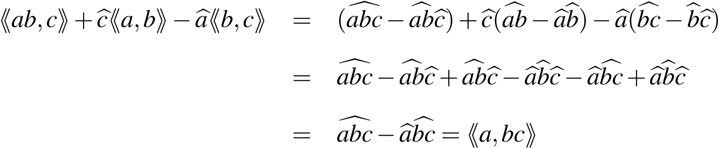
ii. If *c* is independent of *a, b*, and *ab* then ⟪*b, c* ⟫ = ⟪*ab, c* ⟫ = 0, and we have

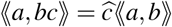

Note that the results of (1) as well can be applied to frequency operation substituting **⟦** ∗, ∗**⟧** for ⟪∗, ∗⟫ and 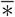 for 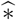.

Recall that for the SIR model the absolute finesses of pathogen strains are independent from each other. With this assumption, the covariance between fitness of two strains will be zero, i.e, **⟦**ŵ _*i*_,ŵ _*j*_**⟧** = 0 and ⟪*w*_*i*_, *w*_*j*_ ⟫ = 0.

**The third term of the Equation** (2) is expanded as a sum of two other terms in Equation (38). Because of the stochastic independency, the covariance between the fitness of two different strains becomes zero and therefore the second term of the Equation (38) becomes zero and we obtain:

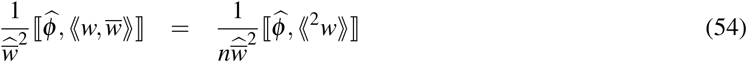

**The fourth term of the Equation** (2) is expanded as a sum two other terms in Equation (39). By part (ii) of Theorem (1), the second term of the Equation (39) become zero and the first term simplified as follows:

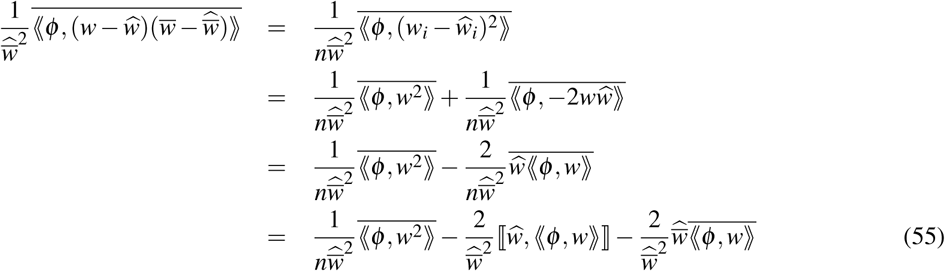

Therefore, using the results of Equations (54) and (55) we simplify Equation (2) as follows:

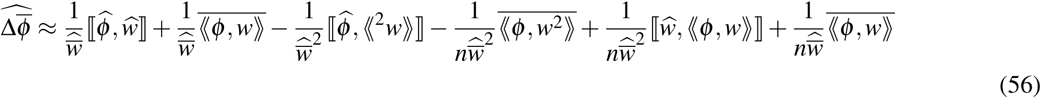

Now we try to obtain each term of the Equation (6). Here, we use Equation (3) and similar equations will be obtained using Equation (4).

Using Equation (5), we expand the **first term on the right-hand side** of the Equation (3) as:

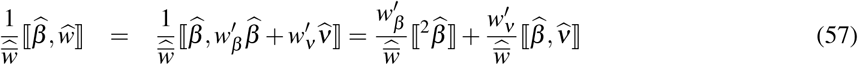

The **second term on the right-hand side** of the Equation (3) can be expanded as:

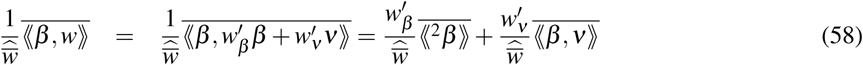

The **third term** of the Equation (3) is expanded as:

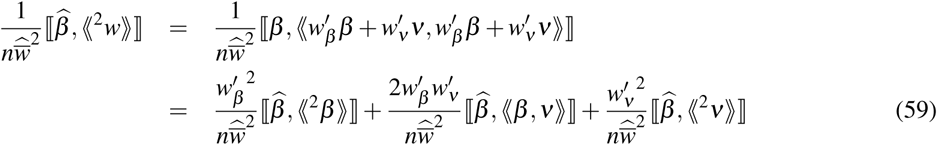

The **fourth term on the right-hand side** of the Equation (3) is expanded below:

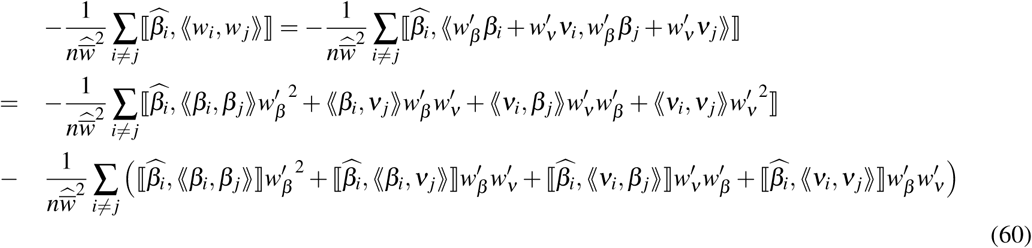

The **fifth term on the right-hand side** of the Equation (3) can be expanded as follows:

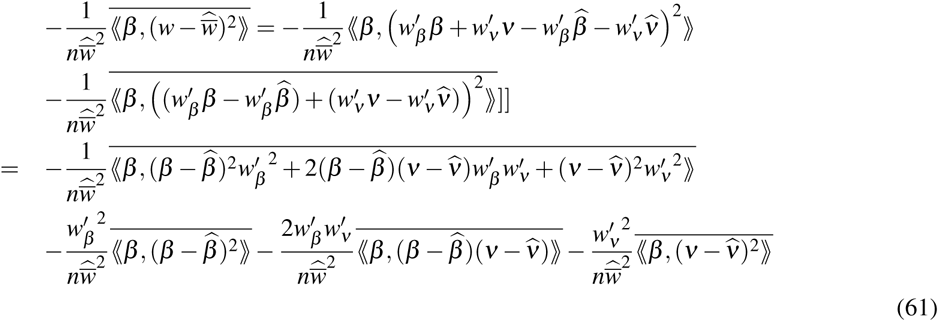

The **sixth term on the right-hand side** of the Equation (3) can be expanded as follows:

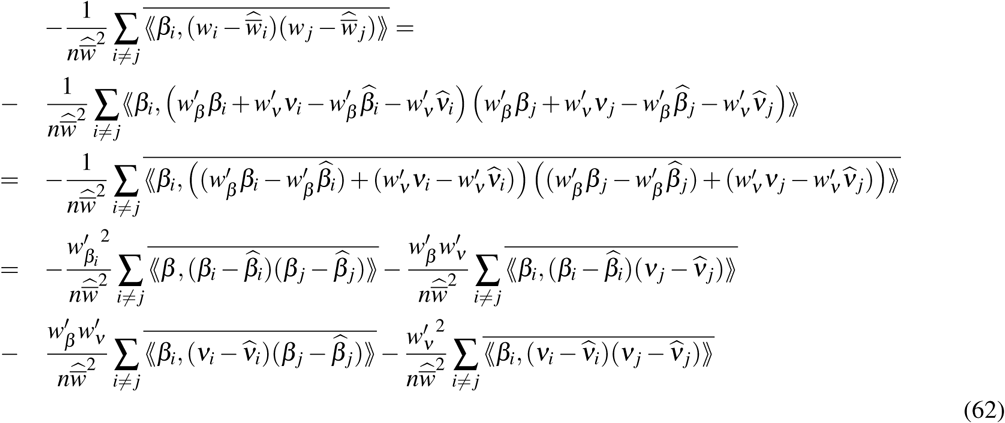

We expand the **seventh term on the right-hand side** of the Equation (3) in two steps:

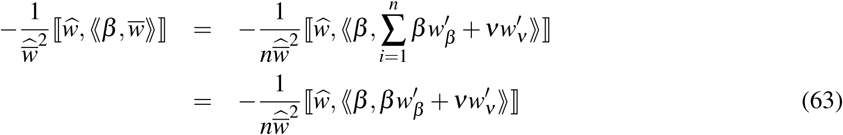

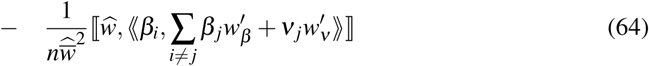

Equation (63) can be further expanded as follows:

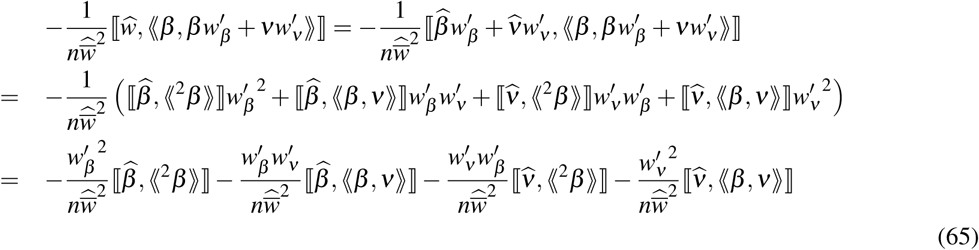

We expand Equation (64) as follows:

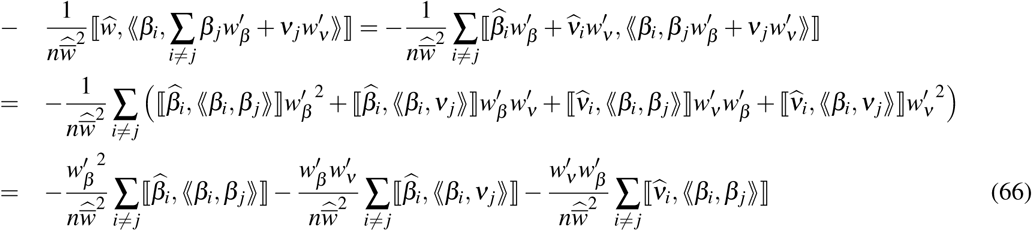

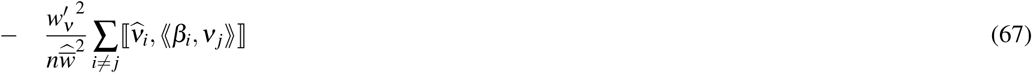

Finally, the **eights term on the right-hand side** of the Equation (3) is expanded below:

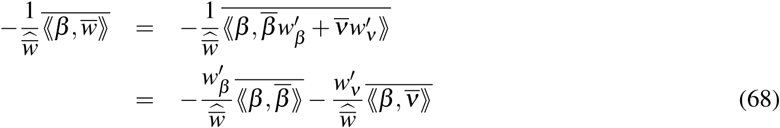

## Appendix 2

From model (22), we derive the equation for fitness of pathogen strain *i* as:

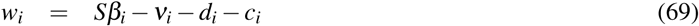

Note that in SIR model the pathogen strains are stochastically independent in fitness. By substituting Equation (69) in Equations (3) and (4) we derive Equation (24) (see Supplementary Materials). We present Equation (24) with the tensor notations. For the strain *i*, we define the random variables 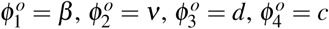. The covariance terms on the right-hand sides of Equations (24) can be grouped into 4 × 4 degree 2 and 4 × 4 × 4 degree 3 tensors (note that a tensor of degree 2 is just a matrix). The elements of each tensor can be shown as follows:

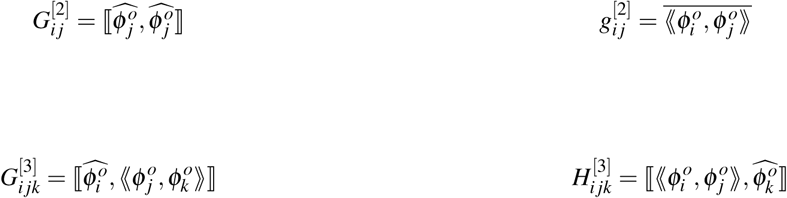

where *i* = 1…4, *j* = 1…4, *k* = 1 4. Also we obtain:

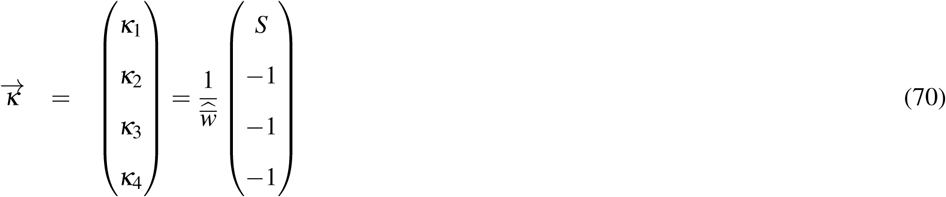

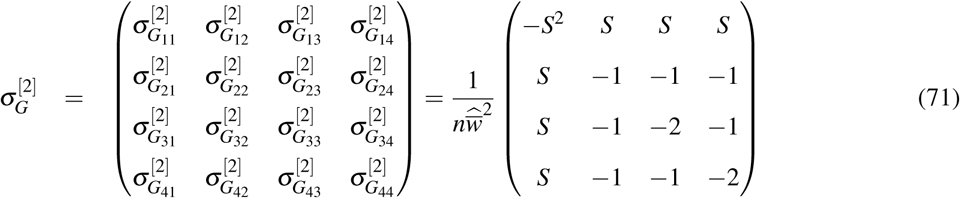

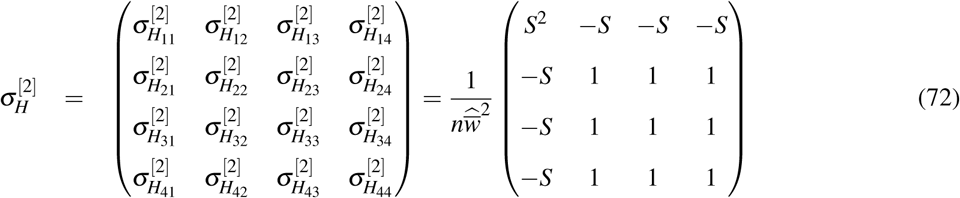

The tensor inner products on the right-hand side of Equation (24) is 4 by 1 vector and can be written as follows:

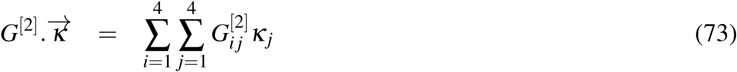

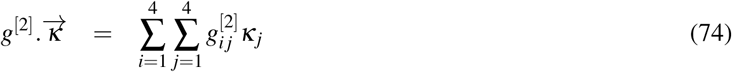

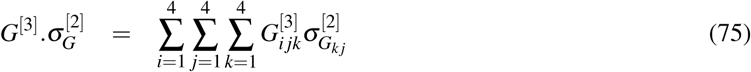

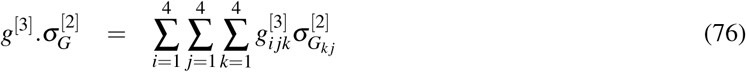

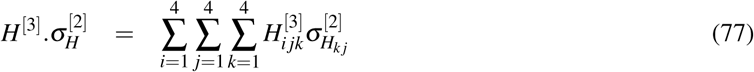

Now we try to derive the Equation (56) for each phenotype *β, ν, d*, and *c* where *w* is function of those variables and obtained from Equation (69). Here we only show the detailed derivation of 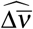 and others can be obtained similiarly.

Using Equation (69), we expand the first term on the right-hand side of the Equation (56) as:

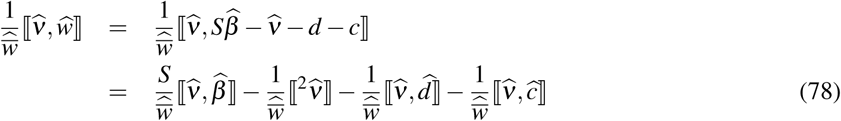

The second term on the right-hand side of the Equation (56) can be expanded as:

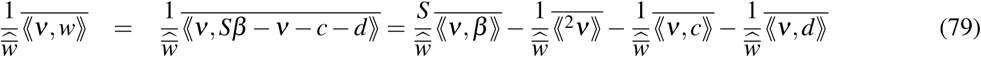

The third term of the Equation (56) is expanded as:

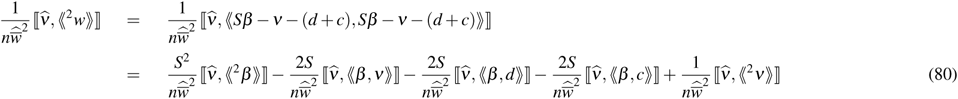

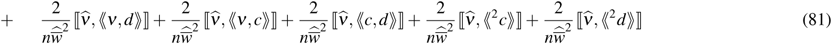

The fourth term on the right-hand side of the Equation (56) is derived below:

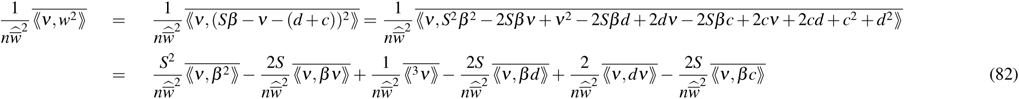

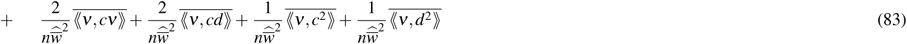

Also, by substituting *ν* in *ϕ* and using Equation (69), the fifth and sixth terms on the right-hand side of the Equation (56) are expanded as follows:

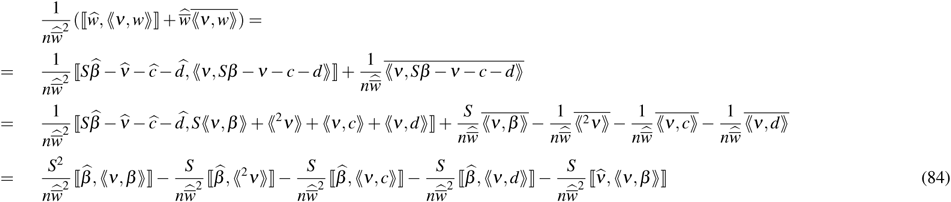

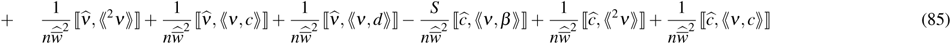

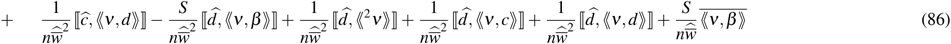

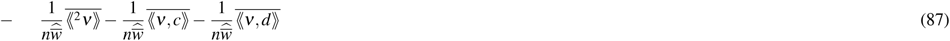

The following equation shows the change in the average mean phenotype of the virulence over one generation. To track where each term comes from, we separate the corresponding terms by writing the equation numbers over them:

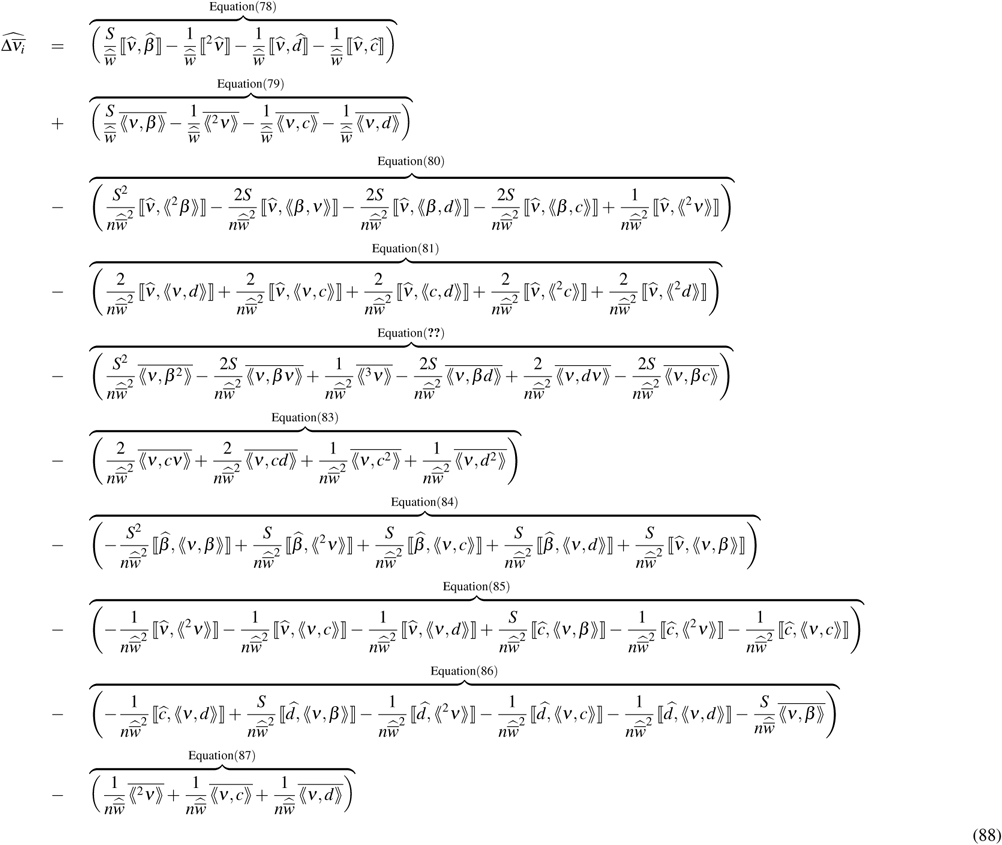

Note that *c* and *d* enter into Equation (69) in the same way as does *ν*. We can thus derive equation 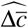and 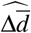 by substituting. Because *β* enters Equation (69) as *Sβ*, the equation for 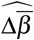 is structured a bit differently. We can use the same approach to derive the equations for the change in the average mean phenotype of the transmission over one generation:

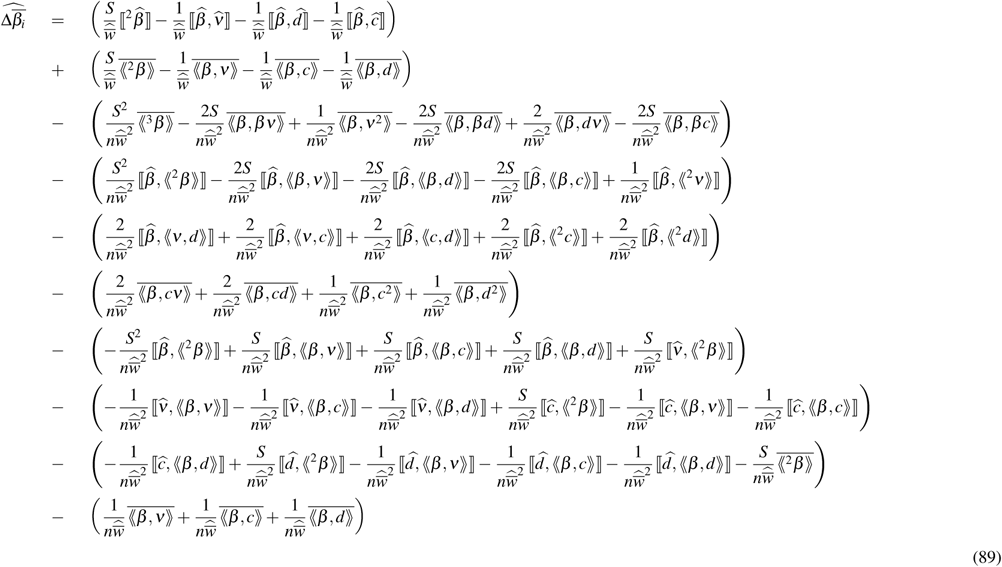

## References

[1] J. H. Gillespie, Natural selection for variances in offspring numbers: a new evolutionary principle, The American Naturalist 111 (1977) 1010–1014.

[2] S. H. Rice, A stochastic version of the price equation reveals the interplay of deterministic and stochastic processes in evolution, BMC Evol Biol 8 (2008) 1.

[3] S. Proulx, Sources of stochasticity in models of sex allocation in spatially structured populations, Journal of evolutionary biology 17 (2004) 924–930.

[4] M. J. Keeling, K. T. Eames, Networks and epidemic models, J R SOC INTERFACE 2 (2005) 295–307.

[5] H. McCallum, N. Barlow, J. Hone, How should pathogen transmission be modelled?, TRENDS ECOL EVOL 16 (2001) 295–300.

[6] J. O. Lloyd-Smith, S. J. Schreiber, P. E. Kopp, W. M. Getz, Superspreading and the effect of individual variation on disease emergence, Nature 438 (2005) 355.

[7] N. Ferrari, I. M. Cattadori, J. Nespereira, A. Rizzoli, P. J. Hudson, The role of host sex in parasite dynamics: field experiments on the yellow-necked mouse apodemus flavicollis, ECOL LETT 7 (2004) 88–94.

[8] A. M. Kilpatrick, P. Daszak, M. J. Jones, P. P. Marra, L. D. Kramer, Host heterogeneity dominates west nile virus transmission, P ROY SOC LOND B BIO 273 (2006) 2327–2333.

[9] S. H. Paull, S. Song, K. M. McClure, L. C. Sackett, A. M. Kilpatrick, P. T. Johnson, From superspreaders to disease hotspots: linking transmission across hosts and space, FRONT ECOL ENVIRON 10 (2012) 75–82.

[10] J. Wolinska, K. C. King, Environment can alter selection in host–parasite interactions, TRENDS PARASITOL 25 (2009) 236–244.

[11] M. E. Woolhouse, C. Dye, J.-F. Etard, T. Smith, J. Charlwood, G. Garnett, P. Hagan, J. Hii, P. Ndhlovu, R. Quinnell, et al., Heterogeneities in the transmission of infectious agents: implications for the design of control programs, P NATL ACAD SCI USA 94 (1997) 338–342.

[12] P. Brodin, M. M. Davis, Human immune system variation, NAT REV IMMUNOL 17 (2017) 21–29.

[13] P. F. Vale, A. J. Wilson, A. Best, M. Boots, T. J. Little, Epidemiological, evolutionary, and coevolutionary implications of context-dependent parasitism, AM NAT 177 (2011) 510–521.

[14] T. Little, P. Vale, M. Choicy, Host nutrition alters the variance in parasite transmission potential, BIOLOGY LETT 9 (2013). doi:10.1098/rsbl.2012.1145.

[15] S. E. Mitchell, E. S. Rogers, T. J. Little, A. F. Read, Host-parasite and genotype-by-environment interactions: temperature modifies potential for selection by a sterilizing pathogen, Evolution 59 (2005) 70–80.

[16] J. W. Mellors, A. Munoz, J. V. Giorgi, J. B. Margolick, C. J. Tassoni, P. Gupta, L. A. Kingsley, J. A. Todd, A. J. Saah, R. Detels, et al., Plasma viral load and cd4+ lymphocytes as prognostic markers of hiv-1 infection, ANN INTERN MED 126 (1997) 946–954.

[17] S. H. Rice, The expected value of the ratio of correlated random variables, unpublished note (2009).

[18] G. R. Price, Selection and covariance., Nature 227 (1970) 520–521.

[19] R. Lande, S. J. Arnold, The measurement of selection on correlated characters, Evolution (1983) 1210–1226.

[20] R. Lande, Natural selection and random genetic drift in phenotypic evolution, Evolution (1976) 314–334.

[21] T. Day, S. Gandon, Insights from price’s equation into evolutionary epidemiology, Disease Evolution: Models, Concepts, and Data Analysis (eds Feng, Zhilan, Dieckmann, Ulf, Levin, Simon A.) 71 (2006) 23–44.

[22] T. Day, S. Gandon, Applying population-genetic models in theoretical evolutionary epidemiology, Ecol. Lett. 10 (2007) 876–888.

[23] J. Harries, Amoebiasis: a review., J ROY SOC MED 75 (1982) 190.

[24] S. Powell, I. MacLeod, A. Wilmot, R. Elsdon-Dew, Metronidazole in amoebic dysentery and amoebic liver abscess, The Lancet 288 (1966) 1329–1331.

[25] R. M. Anderson, R. May, Coevolution of hosts and parasites, Parasitology 85 (1982) 411–426.

[26] S. Alizon, A. Hurford, N. Mideo, M. Van Baalen, Virulence evolution and the trade-off hypothesis: history, current state of affairs and the future, J Evol Biol 22 (2009) 245–259.

[27] H. C. Leggett, C. K. Cornwallis, A. Buckling, S. A. West, Growth rate, transmission mode and virulence in human pathogens, Phil. Trans. R. Soc. B 372 (2017) 20160094.

[28] C. J. Krebs, Why ecology matters, University of Chicago Press, 2016.

[29] E. Papalexi, R. Satija, Single-cell RNA sequencing to explore immune cell heterogeneity, NAT REV IM-MUNOL 18 (2017) 35–45. doi:10.1038/nri.2017.76.

[30] D. Duneau, J.-B. Ferdy, J. Revah, H. Kondolf, G. A. Ortiz, B. P. Lazzaro, N. Buchon, Stochastic variation in the initial phase of bacterial infection predicts the probability of survival in d. melanogaster, eLife 6 (2017). doi:10.7554/elife.28298.

[31] E. Allen, Modeling with Itô stochastic differential equations, volume 22, Springer Science & Business Media, 2007.

